# Long non-coding RNAs as a novel source of beta cell autoantigens in type 1 diabetes

**DOI:** 10.64898/2025.12.01.691527

**Authors:** Jon Mentxaka-Salgado, Aїsha Callebaut, Koldo Garcia-Etxebarria, Leire Bergara-Muguruza, Izei Pascual-González, Anthony R. Jones, Kristen T. Li, Henar Rojas-Márquez, Ainara Castellanos-Rubio, Eddie A. James, Izortze Santin

## Abstract

**Aims/hypothesis:** Genome-wide association studies increasingly highlight the role of non-coding regions in complex diseases such as type 1 diabetes, pointing, in particular, to long non-coding RNAs (lncRNAs) as potentially active molecular players. Emerging evidence suggests that lncRNAs may encode (small) peptides with the potential to modulate cellular processes, including immune responses. In this study, we have investigated whether these (micro)peptides translated from lncRNAs can act as neoantigens capable of activating autoreactive CD4^⁺^T cells in individuals with type 1 diabetes.

**Methods:** We have integrated transcriptomic, proteomic and *in silico* data to identify a subset of lncRNAs with genuine peptide-coding potential in basal or inflamed-condition beta cells. We further assessed the suitability of those candidates to encode (micro)peptides by sequence analysis and *in vitro* experimentation techniques. Finally, we have predicted HLA-II epitopes within the micropeptide candidates and evaluated their immunogenicity by analyzing T cell activation responses in peripheral blood mononuclear cells from HLA matching individuals with type 1 diabetes.

**Results:** By merging ribosome-bound RNA sequencing with nascent peptide mass spectrometry, we identified a total of 30 lncRNAs with potential coding capacity. We verified the peptide-coding ORF translation for three lncRNA candidates: *UXT-AS1*, *RAPGEF4-AS1* and *ENSG00000227066*. After predicting potential type 1 diabetes risk-associated *HLA-DRB1*03:01* and *HLA-DRB1*04:01*-binding epitopes within translated lncRNA (micro)peptides, we identified several peptides that elicited CD4^+^ T cell activation. Furthermore, several epitopes elicited T cell activation in multiple donors with type 1 diabetes. T cell lines were isolated and studied to confirm responses restricted by type 1 diabetes risk HLA alleles.

**Conclusions:** These results reveal a novel class of immunogenic (micro)peptides derived from lncRNAs, supporting their potential role in the autoimmune response that underlies type 1 diabetes. Our findings open new perspectives on the contribution of non-coding genomic elements to autoimmunity and highlight the need to further investigate lncRNA-encoded (micro)peptides as possible targets for future immunotherapies.

**RESEARCH IN CONTEXT:** *What is already known about this subject?:*

- Short proteins can sometimes be translated from lncRNAs.
- Type 1 diabetes neoepitopes arising from non-native forms of proteins and their post translational modifications have been described.

*What is the key question?:*

- Can lncRNA-encoded peptides be recognized by the human immune system and contribute to type 1 diabetes development?

*What are the new findings?:*

- We have identified three candidate lncRNAs with peptide coding capacity.
- Epitopes derived from these peptides drive T cell activation in PBMCs of individuals with type 1 diabetes.
- Presentation of some of these peptides are restricted to type 1 diabetes risk-associated HLA-DRB1*03:01 and *04:01 molecules.

*How might this impact on clinical practice in the foreseeable future?:*

- Discovery of lncRNA-derived peptide reactive T cells might serve as a diagnostic or stratification indicator in the early stages of disease development, aiding the development of more personalized treatments and unlocking a new array of therapeutic targets.

## INTRODUCTION

Type 1 diabetes is a disease in which pancreatic beta cells are destroyed by immune cells, leading to a dependence on exogenous insulin [1]. Before the clinical onset of type 1 diabetes, there is a breakdown of immune self-tolerance, indicated by the accumulation of islet autoantibodies and appearance of autoreactive T cells [2, 3].

Although the autoimmune etiology of type 1 diabetes is undisputed, it is increasingly clear that the beta cell and its dysfunction are key determinants in the immunopathology of the disease [4–6]. Pancreatic beta cells are selectively targeted by infiltrating cells, suggesting inherent features that are specific to beta cells. Notably, CD4^+^ and CD8^+^ T cell populations have been shown to decline with beta cell loss, suggesting an antigen-driven process [7, 8].

Type 1 diabetes is strongly associated with susceptible HLA class II haplotypes, and these molecules dictate the peptide repertoire that is presented to T cells [1, 9]. Hence, presentation of key peptides derived from beta cell antigens by HLA class II molecules must drive specific and targeted autoimmune attack. Thus, identification of relevant antigens and epitopes in type 1 diabetes is a prominent research field [10, 11]. Neoepitopes that occur from post translational modifications of beta cell antigens [12] arise as a result of endoplasmic reticulum (ER) stress, induced by inflammatory responses and/or reactive oxygen species [12–14] and are likely to be underrepresented in the thymus and healthy tissue, evading tolerance mechanisms that limit T cell reactivity [14, 15]. In fact, many classes of neoepitopes have been described in type 1 diabetes, arising from peptide fusions, citrullination, deamidation alternative splicing or even defective ribosomal products (DRiPs) [6, 16].

Still, some classes of disease-relevant epitopes remain largely unexplored. Modern ribosome profiling technologies challenge the traditional view of which genome elements could be translated, as multiple studies report ribosomal footprints outside of annotated protein coding regions [17, 18]. Especially interesting are the lncRNAs, transcripts longer than 200 nucleotides with little to no coding potential which have been linked to regulatory roles in many diseases, and which show differential expression upon pro-inflammatory stimuli [19, 20]. lncRNAs regulate gene expression at multiple levels, including epigenetic modification, transcription, and post-transcriptional processing [21]. Furthermore, several lncRNAs have been shown to encode (micro)peptides from (short) open reading frames (sORFs) within their sequences [22, 23], even in a type 1 diabetes relevant context [24]. This paradigm shift could effectively broaden HLA class II peptide repertoire available for T cell recognition in type 1 diabetes. (Micro)peptides derived from unconventional open reading frames in ribosome-associated lncRNAs, that are upregulated in response to inflammatory stimuli, may constitute a novel source of autoantigens, ultimately contributing to beta cell demise.

Here we report the discovery of several (micro)peptide epitopes derived from translation products of lncRNAs that are recognized by T cells in individuals with type 1 diabetes. Our observations suggest that CD4^+^ T cells that recognize (micro)peptides derived from lncRNAs are a relevant part of the autoreactive repertoire.

## METHODS

### Cell culture

EndoC-βH1 cells (Human Cell Design) were cultured as recommended by the manufacturer. DMEM media containing 2% FBS, 5.6 mmol/l glucose, 50 μmol/l 2-mercaptoethanol (Bio-Rad), 10 mmol/l nicotinamide (Calbiochem), 5.5 μg/ml transferrin and 6.7 ng/ml selenite (Sigma-Aldrich) was used for transfection protocols.

HEK293 cells (CRL-1573) from the American Type Culture Collection (ATCC) were cultured in DMEM supplemented with 10% FBS (Gibco), 100 units/ml penicillin and 100 μg/ml streptomycin (Lonza). Antibiotics was omitted for transfection protocols.

### Intracellular PIC transfection

The synthetic viral dsRNA mimic polyinosinic:polycytidylic acid (PIC) (InvivoGen) was transfected into EndoC-βH1 cells using Lipofectamine 2000 (Invitrogen) at a final concentration of 0.25 μg/ml.

### Ribosome associated RNA sequencing

EndoC-βH1 cells were transfected with PIC for 16 h, and ribosome-associated RNA was purified using the AHARIBO kit (Immagina Biotechnology) following the manufacturer’s instructions. Briefly, cells were methionine-depleted and Azidohomoalanine (AHA) was added to block peptide elongation. Lysis buffer was supplemented with 1% NP40 to improve EndoC-βH1 cell lysis. Ribosomal complexes were purified by cell lysis followed by magnetic AHA affinity pull down. RNA associated to the ribocomplexes was extracted by a Phenol:Cholorofom:Isoamyl alcohol extraction. Sequencing libraries were prepared using “TruSeq Stranded Total RNA with Ribo-Zero Globin” kit (Illumina Inc.) and “TruSeq RNA CD Index Plate” (Illumina Inc.) following “TruSeq Stranded Total RNA Sample Prep-guide”. Sequencing was carried out in a NovaSeq 6000 sequencer. Trimmomatic v 0.39 [25] with default settings used to trim the sequence reads. HISAT 2 [26] was used to align reads using as reference Human Genome assembly hg38 (available from the developer site). Transcripts and their abundances were calculated using Stringtie [27], and transcript counts calculated using htseq-count [28]. Biotypes of the transcripts were annotated using Ensembl 103 release. For differential expression, transcripts with more than 10 counts in at least 4 samples were used. Transcript expression was compared between basal condition and upon intracellular PIC exposure using edgeR package [29] of R language, using the upper quartile method for normalization and sample pairing. enrichR [30] was used to perform gene-set enrichment analyses. Additional statistics analyses and graphs were made using ShinyGO and GraphBio.

### Nascent peptide mass spectrometry

Newly synthesized peptides were purified from actively translating ribosomes following the AHARIBO Protein procedure (Immagina Biotechnology)[31]. Briefly, after AHA treatment and cell lysis, lysates were incubated with AHA-capturing magnetic beads for one hour. Beads were then washed with a urea-containing solution and nascent peptides-bound beads resuspended in water.

Digestion of the nascent peptides was performed with trypsin and lys-C enzymes[31]. Peptides were washed with Sep-Pak C18 1cc Vac Cartridges (Waters) and liquid chromatography was performed on an EASY nLC-1200 machine (Thermo Scientific) with a 90-minute gradient. MS/MS fractionation was performed in an Exploris 480 (Thermo Scientific). Protein identification was performed using the Protein Discoverer 2.2 software (Thermo Scientific) against the human proteome database UniProtKB plus a custom-built database containing potential translations of the ORFs of interest. Parameters were set to 2 missed cleavage allowance, post translational modifications (carbamidomethylation, oxidation and acetylation) and a false discovery rate (FDR) of 1%.

### Gene expression measurement

Total RNA extraction of EndoC-βH1 was performed using the NucleoSpin RNA kit (Macherey-Nagel), following manufacturer’s instructions. Quantitative PCR was performed using iTaq Universal One-Step RT-qPCR Kit (BioRad) in a C1000 Touch Thermal cycler (Bio-Rad) with specific primers (**ESM Table 1**).

**Table 1.** Summary of the identified lncRNA-derived (micro)peptides by merging ribosome-associated RNAseq and nascent peptide MS data.

| **Gene** | **Transcript** | **Putative**<br>**peptide length**<br>**(aa)** | **Experimental tryptic peptide** |
| --- | --- | --- | --- |
| *UXT-AS1* | *ENST00000669654* | 90 | LGDIAAAASVQR |
| *RAPGEF4-AS1* | *ENST00000435328* | 137 | DQVFFISLYPLISR |
| *ENSG00000227066* | *ENST00000431027* | 214 | TVNIYTDSR |
| *LINC03000* | *ENST00000669047* | 106 | EDTGELR |
| *LINC01572* | *ENST00000664779* | 57 | ISWLALYCDK |
| *ENSG00000285517* | *ENST00000648050* | 138 | IYLFLR |
| *GARS1-DT* | *ENST00000581665* | 105 | LIFPAGNELK |
| *TMCC1-DT* | *ENST00000671446* | 111 | MSLMLR<br>LSQMAEENLRK |
| *ENSG00000272970* | *ENST00000610001* | 30 | MIAEGHTAWLQASR |
| *RIPK2-DT* | *ENST00000691446* | 30 | MINPVIVEQVQVFNR |
| *LINC01588* | *ENST00000528300* | 46 | MSFPHLR |
| *ST8SIA5-DT* | *ENST00000661793* | 53 | SPLRPVGR |
| *EFCAB10-AS1* | *ENST00000609827* | 59 | SWQHFFGMLK |
| *GMDS-DT* | *ENST00000655727* | 129 | VCVHVCACMWTCVYVCMCVCK |
| *LINC00506* | *ENST00000628435* | 52 | VLCQGINSR |
| *ENSG00000228566* | *ENST00000660795* | 34 | VTFLTIR |
| *APPBP2-DT* | *ENST00000691821* | 60 | ASLPSGPGVEWSGVDR |
| *RORA-AS1* | *ENST00000558140* | 29 | IHIVCAMVLSQLR |
| *PCBP1-AS1* | *ENST00000630975* | 343 | ILGVWDTAVSLGK |
| *PKN2-AS1* | *ENST00000425750* | 51 | ISMIPNVTR |
| *ENSG00000223930* | *ENST00000411727* | 63 | LGAQYQPPR |
| *LINC00870* | *ENST00000664354* | 31 | MARMPR |
| *ENSG00000232732* | *ENST00000596619* | 33 | MEHAQQQGRPQSR |
| *ENSG00000260337* | *ENST00000658288* | 43 | MPCAYPVPPVGTAPPFTSMPR |
| *EOLA2-DT* | *ENST00000444489* | 37 | NLELPSR |
| *ENSG00000225087* | *ENST00000654386* | 41 | NPDYLHFKCQMKER |
| *ENSG00000273064* | *ENST00000608069* | 101 | RPICNMNWQSQEPGCK |
| *LINC02577* | *ENST00000650196* | 26 | SLNLLSVSWRPR |
| *ENSG00000270792* | *ENST00000603985* | 26 | TPWITYNLVGPR |
| *ENSG00000224090* | *ENST00000441450* | 65 | TQEQT LGHGDSR |

Expression of lncRNAs of interest was measured in a commercial panel of human tissues (Human total RNA master panel II, Clontech).

All qPCR measurements were performed in duplicates, and expression levels were assessed using the 2^−ΔΔCt^ method.

### Determination of lncRNA subcellular localization

To assess cellular localization, cytoplasmic and nuclear RNA fractions of EndoC-βH1 cells were purified using Cytoplasmic and Nuclear RNA Purification Kit (Norgen). Expression levels of *RPLP0* (as a cytoplasmic control RNA), *Lnc13* (as a nuclear control RNA)[32] and transcripts of interest were measured by RT-qPCR.

### Candidate ORF overexpression

The sequences of lncRNA candidates of interest were ordered as gBlock gene fragments (IDT). The ORFs followed by a FLAG tag were cloned into pCMV6 mammalian expression plasmids using FseI and KpnI restriction enzymes. Control or overexpression vectors were transfected into HEK293 cells using X-tremeGENE HP (Sigma) following manufacturer’s instructions. Cells were harvested 16, 24, 48 and 72 hours post-transfection. Transfection in the EndoC-βH1 cells was performed using Lipofectamine-2000 following manufacturer’s instructions, and proteins harvested 24 hours post-transfection.

### m^6^A methylation analysis

m^6^A methylation marks in *UXT-AS1, RAPGEF-AS1* and *ENSG00000227066* transcripts were predicted using SRAMP[33]. To confirm m^6^A methylation 3 μg of precleared RNA per sample was fragmented with RNA fragmentation buffer (100 mM Tris, 2 mM MgCl2) for 3 min at 95°C and placed on ice. 20% of RNA was kept as input. 1 μg of m^6^A antibody (Abcam, #ab151230) and control antibody (IgG, Santa Cruz Biotechnologies, Dallas, USA, #sc-2025) were coupled to magnetic beads (ThermoFisher Scientific, MA, USA) in a rotation wheel for 1 h at 4°C. After incubation, beads were washed twice in reaction buffer (150 mM NaCl, 10 mM Tris-HCl, 0.1% NP-40). RNA was added to the antibody-coupled beads and rotated for 2 h at 4°C. Subsequently, beads were washed 3X in reaction buffer, 3X in low salt buffer (50 mM NaCl, 10 mM TrisHCl, and 0.1% NP-40) and 3X in high salt buffer (500 mM NaCl, 10 mM TrisHCl, and 0.1% NP-40). Beads were then resuspended in lysis buffer and RNA was extracted using the PureLink RNA extraction kit (Invitrogen, Carlsbad, USA, #12183016). qPCR of the regions of interest was performed using the primers listed in **ESM Table 1**.

### Western Blot

Cells were lysed in Laemmli sample buffer (0.0625 M Tris-HCl, 2% SDS, 10% glycerol, 0.005% bromophenol blue) supplemented with 5% beta-mercaptoethanol (Bio-Rad) and run in a 14% acrylamide gels. Proteins were transferred to a 0.2 µm pore PVDF membrane in a wet Transfer System (Bio-Rad, 100V for 1h) and blocked in non-fatty 5% milk TBST 1X (0.1% Tween 20 TBS) for 1 hour. Membranes were incubated with anti-FLAG (1:1,000; Sigma-Aldrich, #F3165) or anti-β-tubulin (dilution 1:5,000; Proteintech, #66031) overnight, followed by one-hour incubation with a secondary anti-mouse HRP-conjugated antibody (Dilution 1:10,000; Santa Cruz Biotechnologies, #sc-516102). Proteins were detected in a Chemidoc XRS+ (Bio-Rad) using the chemiluminescent Clarity Max Western ECL substrate (Bio-Rad).

### Immunofluorescence

After transfection, EndoC-bH1 cells were fixed with 4% paraformaldehyde, and permeabilized with 0.3% Triton X-100 (Sigma). The cells were then incubated overnight with the FLAG antibody (1:1000; Sigma-Aldrich). Alexa-Fluor 568 mouse antibody (1:500; ThermoScientific; #A11061) was used for visualization by fluorescence microscopy in a Zeiss LSM880 Fast Airyscan microscope.

### Structural and localization predictions

Peptides of interest were analyzed using AlphaFold[34] for three-dimensional structure prediction and confidence estimation (pLDDT). Physicochemical properties were calculated with ExPASy ProtParam [35], and subcellular localization was inferred using DeepLoc 2.0[36].

### HLA-II competitive peptide binding assay

Putative epitopes from ribosome-associated lncRNA peptides (**ESM Table 2**) were predicted as previously described [37]. Binding affinity was measured by incubating biotinylated B-myoglobulin 137-148 (HLA-DRB1*03:01) or influenza hemagglutinin 306-318 (HLA-DRB1*04:01) in wells coated with recombinant DRB1*03:01/HLA-DRB1*04:01 [37]. Residual biotinylated peptide was detected with europium-streptavidin on a VICTOR Nivo time-resolved fluorometer (Perkin Elmer).

**Table 2.**
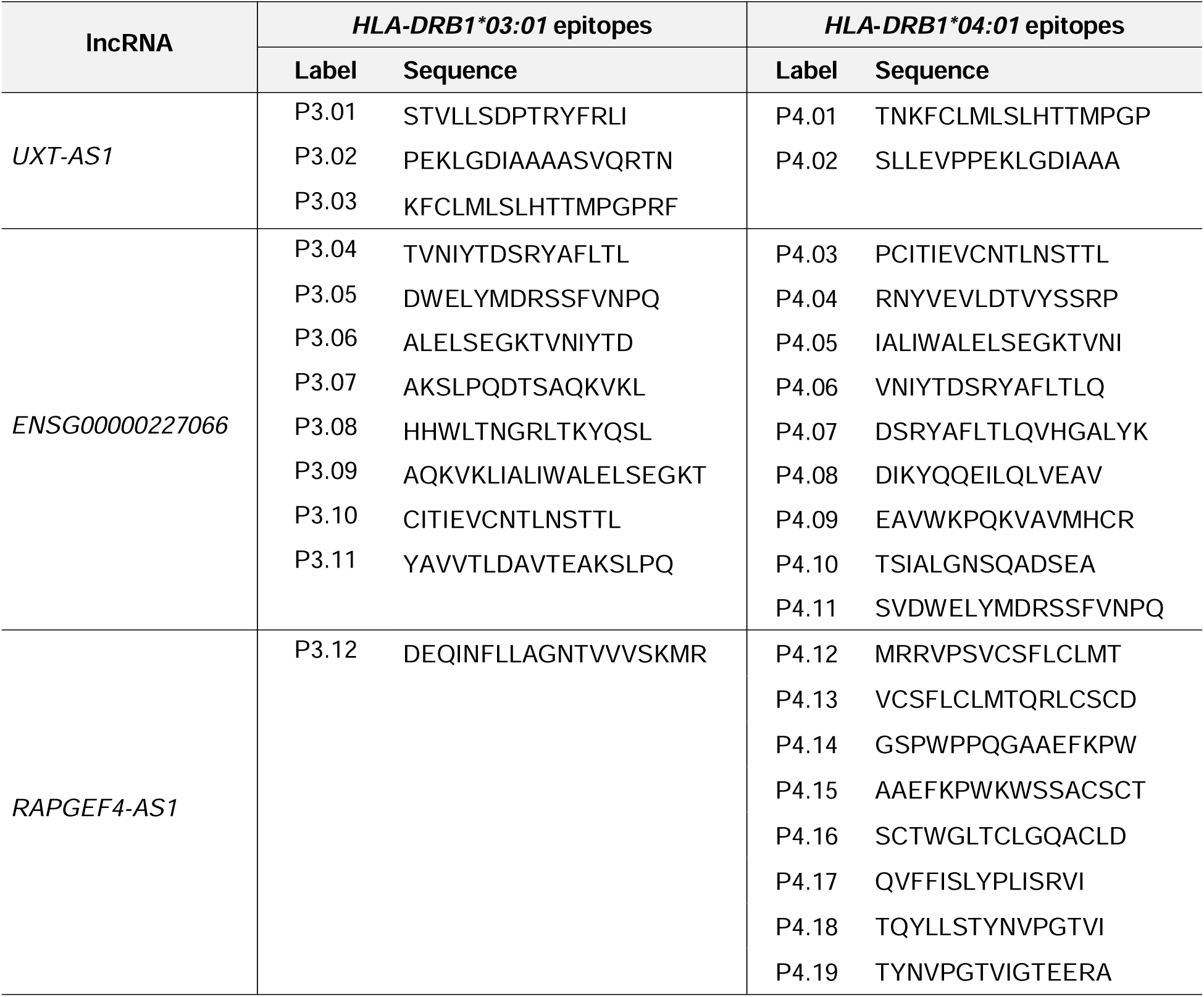
List of the predicted immunogenic epitopes in lncRNA-encoded peptides.

### In vitro expansion of human PBMCs

Frozen PBMCs from individuals with type 1 diabetes recruited with informed consent under a study approved by the institutional review board at the Benaroya Research Institute (**ESM Table 3**) were thawed and resuspended in medium consisting of RPMI 1640 + 10% FBS + 2 mM L-glutamine + 50 U/ml PenStrep (all from Thermo Fisher) supplemented with Benzonase (Sigma). PBMCs were cultured in T Cell Medium consisting of RPMI 1640 + 15% Human serum (in house preparation), 2 mM L-glutamine, and 50 U/ml penicillin-streptomycin and stimulated with peptides for 21 days, adding medium and IL-2 starting on day 7.

**Table 3.**
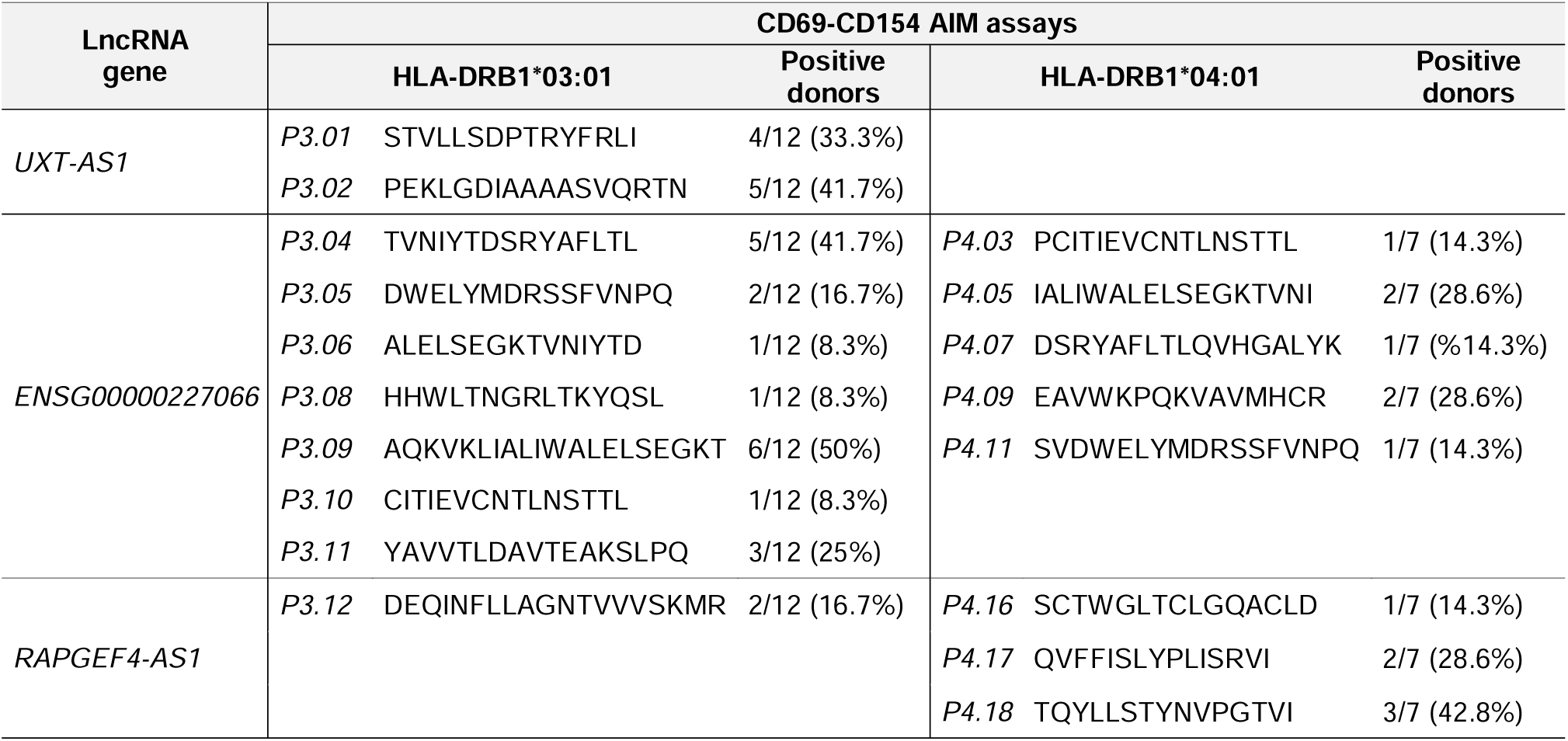
Summary of primary positive responses against HLA-DRB1*03:01 and HLA-DRB1*04:01 peptides in PBMCs from HLA-DRB1-matched individuals with type 1 diabetes. Those peptides which didn’t cause any activation responses in any of the donor PBMCs have been omitted.

### Activation-induced marker (AIM) assays

T cells were stimulated for 5h at 37 °C with peptide (0.1 mg/ml) in the presence of anti-CD40 antibody (2 µg/mL) and stained for 30 minutes at 4 °C in the dark with CD4-BUV395 (BD Biosciences), CD3-PerCP (Biolegend), CD69-APC (Biolegend) and CD154-PE (Miltenyi) antibodies. To obtain lines, CD154^+^, CD69^+^, CD4^+^ T cells were sorted into round bottom wells (4 cells per well) plus 10^5^ irradiated PBMCs and 2 µg/mL phytohemagglutinin (Remel Inc.). Fresh TCM supplemented with human 10 U/ml IL-2 was added every two days.

### T cell line proliferation assays

T cell lines were plated in duplicate with irradiated antigen presenting cells (APCs) expressing one HLA molecule of interest[38]. These included immortalized B cell lines transduced to express HLA-DRB1*03:01- or HLA-DRB1*04:01. T cells were stimulated with 10 mg/mL of each peptide. After incubation for 72 h, cells were pulsed with medium containing ^3^H-thymidine (1 mCi/well). Incorporation was measured 18 h later with a scintillation counter. A stimulation index (SI) was calculated by normalizing the ^3^H-thymidine incorporation with that of non-stimulated wells.

### Statistics

Experimental data are displayed as means±SEM. Comparisons between groups were performed by Student’s t test or one-way ANOVA followed by correction for multiple comparisons (as recommended by GraphPad Prism v8.0.1). p-values below 0.05 were considered significant.

## RESULTS

### lncRNAs harbor ORFs with coding potential

To identify lncRNA candidates that are potentially translated into (micro)peptides, we combined RNA expression and mass spectrometry analysis of basal and PIC-treated EndoC-βH1 cells (**Fig. 1a).** RNA sequencing of ribosome-associated RNA identified 33,925 and 38,635 unique ribosome-associated transcripts in basal and PIC conditions respectively, of which lncRNAs represented 27%. We performed differential gene expression analysis of lncRNA and protein coding genes that had 10 counts or more in at least 4 samples (12,357 genes and 3,245 lncRNAs were included). Consistent with previous studies, differential gene expression analysis confirmed that PIC treatment upregulated a plethora of pro-inflammatory and antiviral response genes (i.e. *ISG15*, *MX1*, *OAS1*, and *IRF7*) [19, 39], validating PIC as a suitable viral mimic (**ESM Fig. 1a**). Gene Ontology (GO) enrichment analysis revealed enriched upregulated coding genes in pathways related to innate immune response, antiviral response and type I IFN signaling, among others (**ESM Fig. 1b**). Differential gene expression analysis of the lncRNAs identified a 454 gene subset upregulated in ribosomes in response to PIC (FDR<0.05 and −log2FC≥1.5) (**Fig. 1b**). In contrast, only 3 lncRNAs showed significant downregulation in PIC-treated EndoC-βH1 cells compared to control cells (FDR<0.05 and −logFC≤-1.5). Although most of the detected lncRNA transcripts were not annotated, some had been already described as coding in other tissues. For example, lncRNA *CRNDE* encodes a nuclear 84 amino acid micropeptide that participates in cell proliferation [40], and *Linc-PINT* encodes a micropeptide of 87 amino acids expressed in glioblastoma cells [41]. These results validated the utility of this approach to detect potentially encoding lncRNAs.

**Fig. 1.**
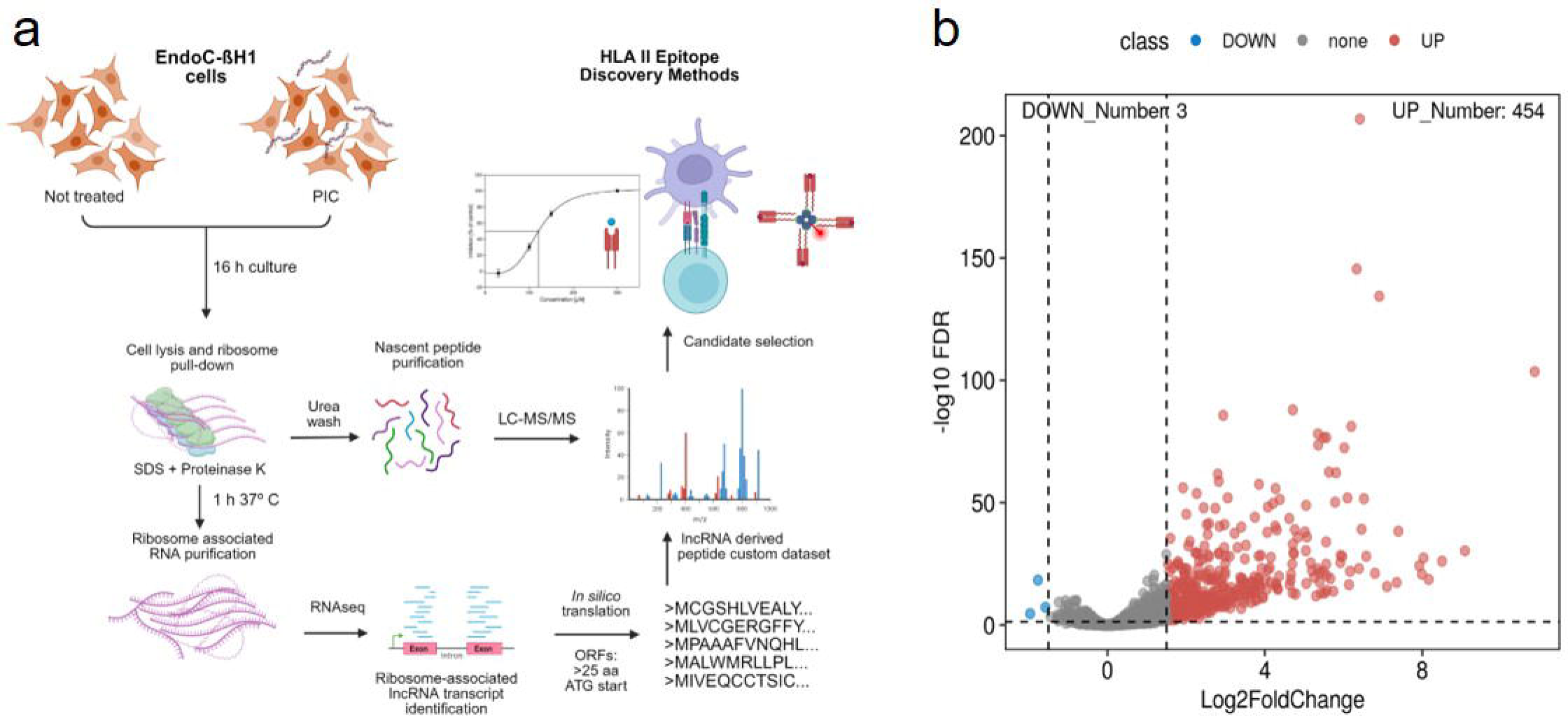
Several lncRNAs are upregulated in ribosomal complexes after PIC stimulation in pancreatic beta cells. a) Pipeline designed for the identification of lncRNA transcripts with coding potential, combining mass spectrometry and RNA sequencing. b) Volcano plot showing the differential expression of the ribosome associated lncRNAs. Downregulated lncRNAs (FDR<0.05 and −logFC≤-1.5) are represented as blue dots and upregulated ones as red dots (FDR<0.05 and - log2FC≥1.5).

Then, we analyzed the nascent peptide translatome purified from active ribosomal complexes by standard LC-MS/MS. Spectra were matched against a custom database containing both the human proteome (UniprotKB release 2023_05), and theoretical translations of ORFs from all lncRNA transcripts detected in the ribosome-associated RNAseq (a total of 96,370 ORFs) (**Fig. 1a**). Only open reading frames starting with the canonical ATG codon and longer than 75 nucleotides were considered. Altogether, by MS analysis of ribosome-bound nascent peptides identified ∼1,900 human proteins with high confidence in each of the 4 samples. We obtained 32 hits that matched theoretical translations of our custom lncRNA ORF database (**Table 1**). The identified 32 tryptic peptides came from 31 unique ORF translations, and only one peptide was identified by two different experimental tryptic peptides (*TMCC1-DT* translation product). Matches were distributed roughly in half between basal and PIC treated samples, indicating no enriching effect of PIC stimulus on the production of lncRNA-derived (micro)peptides. One candidate was dropped from our list, as verification of the ORF by BLAT indicated that it was a translation product of a transcript annotated as coding in GENCODE v47 (Human *APRV1*).

### Stable peptides are encoded from lncRNAs

Next, we checked the ability of candidate lncRNA ORFs (**ESM Table 2**) to encode peptides. Overexpression of 3’ end FLAG-fused ORF constructs in HEK293 cells yielded detectable peptide translation for *RAPGEF4-AS1, UXT-AS1* and *ENSG00000227066* lncRNAs at the expected molecular weights of 17 KDa, 12 KDa and 26 KDa, respectively (**Fig. 2a**). The three lncRNA peptides were detectable at 24 and 48h post-transfection, although expression of the peptide encoded by *ENSG00000227066* (ENS) lncRNA was reduced at 48h, suggesting that this peptide had lower stability. In contrast, the peptide encoded from lncRNA *UXT-AS1* was still detectable after 72h of transfection, suggesting high stability. These results were confirmed in the pancreatic beta-cell line EndoC-βH1 (**Fig. 2b**). Although transfection of the lncRNA ORF-FLAG constructs was less efficient than in HEK293 cells (data not shown), all three (micro)peptides of interest were successfully detected by immunofluorescence using an anti-FLAG antibody.

**Fig. 2.**
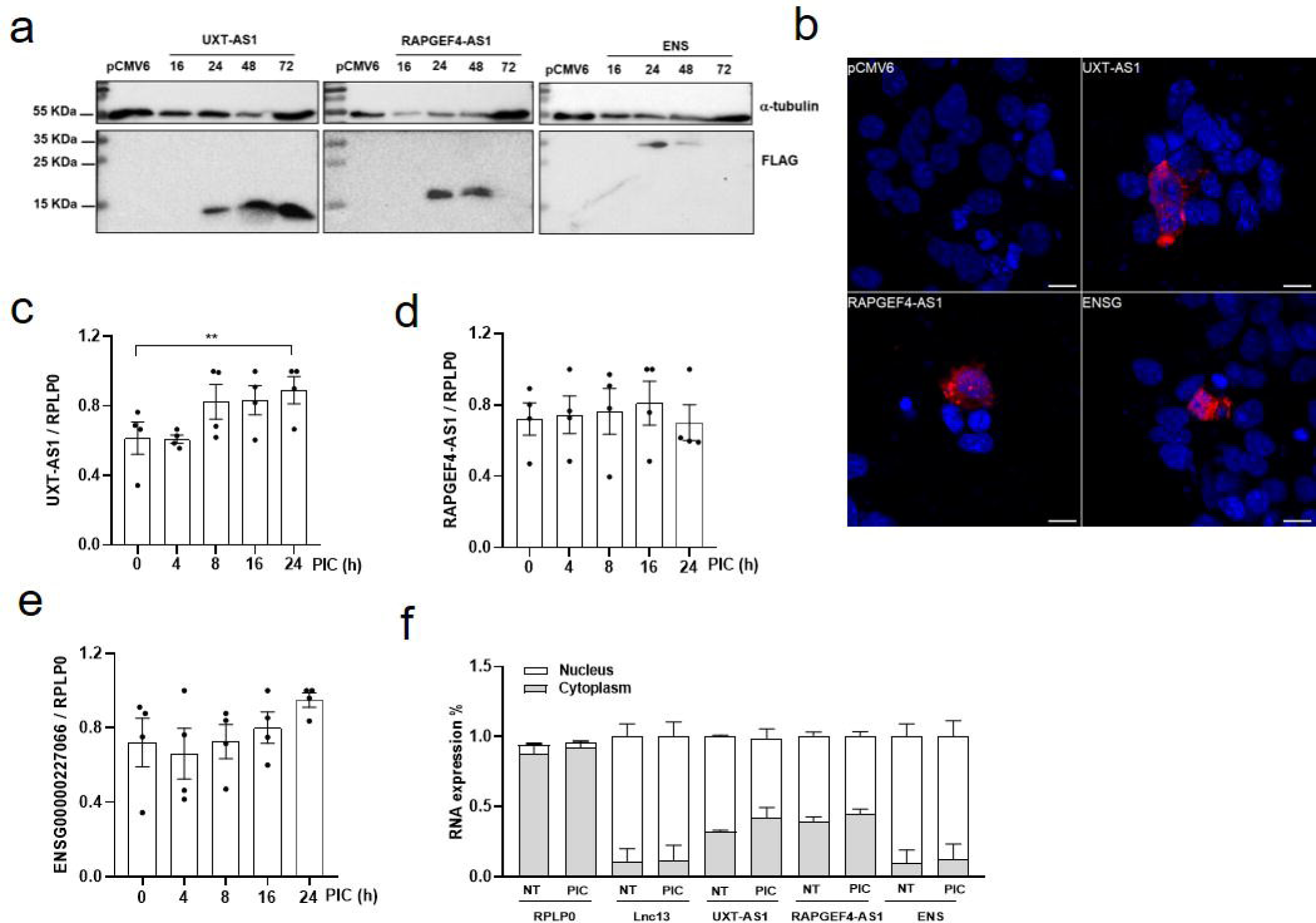
Ribosome-associated lncRNAs encode stable (micro)peptides. a) Cells were transfected with an empty overexpression vector (pCMV6) or vectors containing the ORFs of *UXT-AS1, RAPGEF4-AS1* or *ENSG00000227066* (ENS) fused to a flag tag for 16, 24, 48 or 72h. The expression of the peptides was detected using an anti-FLAg antibody, and α-tubulin was determined as loading control. b) Cells were transfected with an empty overexpression vector (pCMV6) or vectors containing the ORFs of *UXT-AS1, RAPGEF4-AS1* or *ENSG00000227066* (ENS) fused to a flag tag for 24h. Expression of the lncRNA-encoded peptides was detected by immunofluorescence. Scale: 10μm. c-e) EndoC-βH1 cells were left untreated (0h) or exposed to intracellular PIC (0.25 μg/mL) for 4, 8, 16 or 24h. The expression of *UXT-AS1* (c), *RAPGEF4-AS1* (d) and *ENSG00000227066* (e) was determined by qPCR. Results are means±SEM of 4 independent experiments; **p<0.01; Student’s t test. f) EndoC-βH1 cells were left non-transfected (NT) or exposed to PIC (0.25 μg/mL) for 16h (PIC). Relative *UXT-AS1, RAPGEF4-AS1* and *ENSG00000227066* (ENS) b expression was determined in nucleus and cytoplasmic fractions. *RPLP0* and *Lnc13* were determined as controls for the cytoplasmic and nuclear fractions, respectively. Data are expressed as % RNA expression and are means±SEM of 3 independent experiments.

Sequence-based analyses revealed distinct structural characteristics among the three proteins (**ESM Fig. 2**). The peptide encoded by *ENSG00000227066* displayed a well-defined globular domain with high-confidence regions (pLDDT > 90) connected by flexible linkers, consistent with a soluble protein (**ESM Fig. 2a**). DeepLoc predicted dual localization in the cytoplasm and nucleus. The peptide encoded from *UXT-AS1* showed low structural confidence (pLDDT < 50) and contained a hydrophobic N-terminal stretch with six cysteine residues that may allow transient structural stabilization through disulfide bonds (**ESM Fig. 2b**). DeepLoc predicted localization of this peptide to the nucleus and mitochondrion. The *RAPGEF4-AS1*-derived peptide also exhibited low-confidence structural prediction (pLDDT < 50) and was also rich in cysteines. Based on DeepLoc prediction, this peptide was classified as cytoplasmic (**ESM Fig. 2c**).

These lncRNAs had not previously been shown to produce any translated (micro)peptide. *In silico* analysis of the candidates showed that the *ENSG00000227066* transcript was classified as “coding” by the CPC2 algorithm (score: 0.99), predicting the 216 amino acid peptide identified in our mass spectrometry experiments. Additionally, we found ribosome footprint evidence in the TransLnc database supporting ribosomal occupancy of this lncRNA transcript, albeit with different predicted ORF translations. In contrast, the *UXT-AS1* and *RAPGEF4-AS1* transcripts were classified as “non-coding” by CPC2 (scores: <0.5). While some ribosome profiling evidence was annotated in Translnc database for *RAPGEF4-AS1* transcript, no evidence of ribosome footprint was described for *UXT-AS1* lncRNA. None of the lncRNAs presented evolutionary conservation of their nucleotide sequence among vertebrates based on the PhyloCSF algorithm (**ESM Fig.3**).

**Fig. 3.**
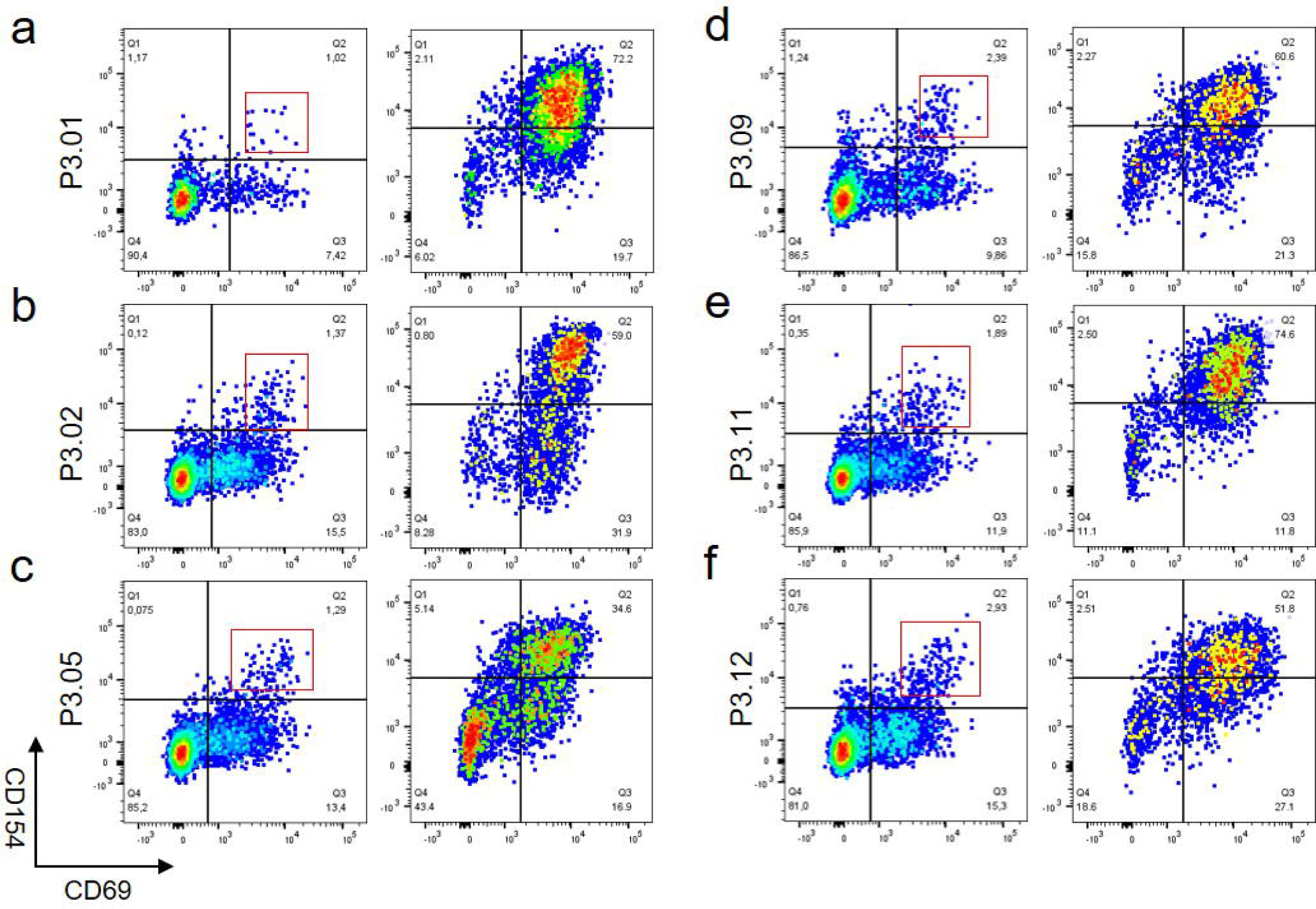
HLA-DRB1*03:01 epitopes predicted from lncRNA derived peptides drive T cell activation. a-f) Each panel shows a representative in vitro primary activation response (left plots) and an illustrative T cell line expanded from the positive populations (right plots) against a specific peptide.

To determine the expression patterns of these peptide-encoding lncRNAs, we first analyzed their expression in a set of human samples, observing that all three lncRNAs are ubiquitously expressed in different human tissues (**ESM Fig.4**). The highest expression of *UXT-AS1* was observed in brain tissue, and the lowest in salivary glands. For *RAPGEF-AS1* lncRNA, the highest expression was detected in the heart, and the lowest in the prostate. For *ENSG00000227066,* the highest expression was observed in the thyroid, and the lowest in muscle. Single cell RNAseq data from Human Pancreas Analysis Program (HPAP) suggests that expression of *UXT-AS1* and *RAPGEF4-AS1* is very low in human pancreatic beta cells (0.099 TPMs and 0.027 TPMs, respectively). Interestingly, the expression of both was slightly upregulated in the pancreatic beta cells of individuals with type 1 diabetes (1.5- and 2.5-fold, respectively). Expression data for *ENSG00000227066* was not available. It is important to highlight that the values reported in HPAP may not reflect absolute expression levels, given that single-cell RNA sequencing lacks adequate sensitivity detection and quantification of low-abundance lncRNAs [42]. We next analyzed the expression of the three lncRNAs in the human EndoC-βH1 cell line in basal condition and after transfection with PIC. Our analysis confirmed that the basal expression of the three lncRNAs was low (**Fig. 2c-e**), however the expression of *UXT-AS1 was* slightly upregulated after 24 h of PIC transfection (**Fig. 2c)**. The expression of *RAPGEF4-AS1* and *ENSG00000227066* did not change upon intracellular PIC exposure (**Fig. 2d-e**).

**Fig. 4.**
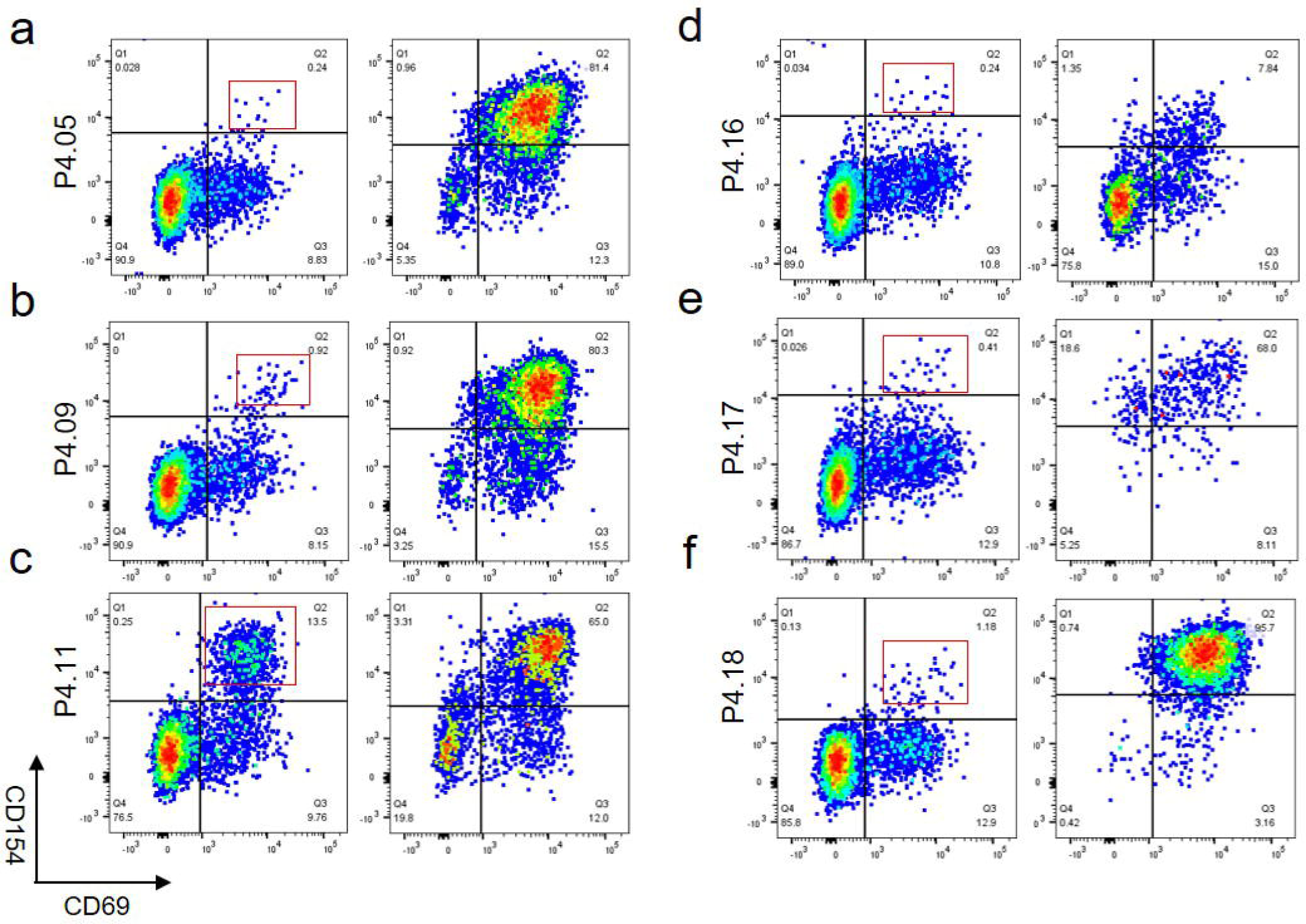
HLA-DRB1*04:01 epitopes predicted from lncRNA derived peptides drive T cell activation. a-f) Each panel shows a representative in vitro primary activation response (left plots) and an illustrative T cell line expanded from the positive populations (right plots) against a specific peptide.

Subsequently, we examined the subcellular localization of the three lncRNAs in EndoC-βH1 cells under basal conditions and upon PIC transfection (**Fig. 2f**). LncRNAs that are prone to be translated are thought to localize in the cytoplasm. *UXT-AS1* and *RAPGEF4-AS1* were equally distributed in the nucleus and the cytoplasm and PIC transfection did not affect their localization. Regarding *ENSG00000227066*, around 85% of the transcript molecules were in the nuclei both at basal and PIC-transfected conditions.

The ability of a lncRNA to encode a peptide can be influenced by several molecular features, including the presence of internal ribosome entry sites (IRES) or epitranscriptomic modifications such as N6-methyladenosine (m^6^A) in the 3’ untranslated region (3’ UTR) [43, 44]. We analyzed these features to determine whether the lncRNAs of interest possess elements that explain their capacity for translation. *In silico* analysis of IRES sequences using SIRES webserver revealed a high confidence IRES in *UXT-AS1*, however, its localization inside the ORF of interest indicated an improbable functional role in initiating translation of the full-length peptide (data not shown). We next examined the presence of m^6^A epitranscriptomic motifs within the three lncRNAs of interest. We performed an *in silico* analysis using an online m^6^A predictor [45] (**ESM Fig. 5**). LncRNAs *UXT-AS1* and *RAPGEF4-AS1* were predicted to harbor a unique m^6^A motif classified as “high confidence” and “very high confidence”, respectively. While the m^6^A motif in *UXT-AS1* was located inside the ORF of interest (**ESM Fig. 5a**), in *RAPGEF4-AS1* transcript it was located in the 3’UTR of the ORF of interest (**ESM Fig. 5b**). On the other hand, *ENSG00000227066* transcript was predicted to harbor five m^6^A motifs classified as “very high confidence”. All were located in the 3’UTR of the ORF of interest (**ESM Fig. 5c**). To confirm the presence of m^6^A modifications, we next performed an m^6^A-RIP using RNA extracted from basal and PIC-transfected EndoC-βH1 cells, followed by qPCR analysis of the regions of interest. Neither *UXT-AS1* nor *RAPGEF4-AS1* showed m^6^A methylation under basal or PIC-transfected conditions (**ESM Fig. 6a-b**). For *ENSG00000227066*, two primer pairs were designed: one targeting the m^6^A site at position 1028 (m^6^A_7) and another targeting the region containing m^6^A sites between positions 1844 and 1930 (m^6^A_11-15). As shown in **ESM Fig. 6c**, no methylation was detected at the m^6^A_7 site. Interestingly, while the m^6^A_11-15 region was unmethylated at basal conditions, it became significantly methylated following intracellular PIC exposure (**ESM Fig. 6d**).

**Fig. 5.**
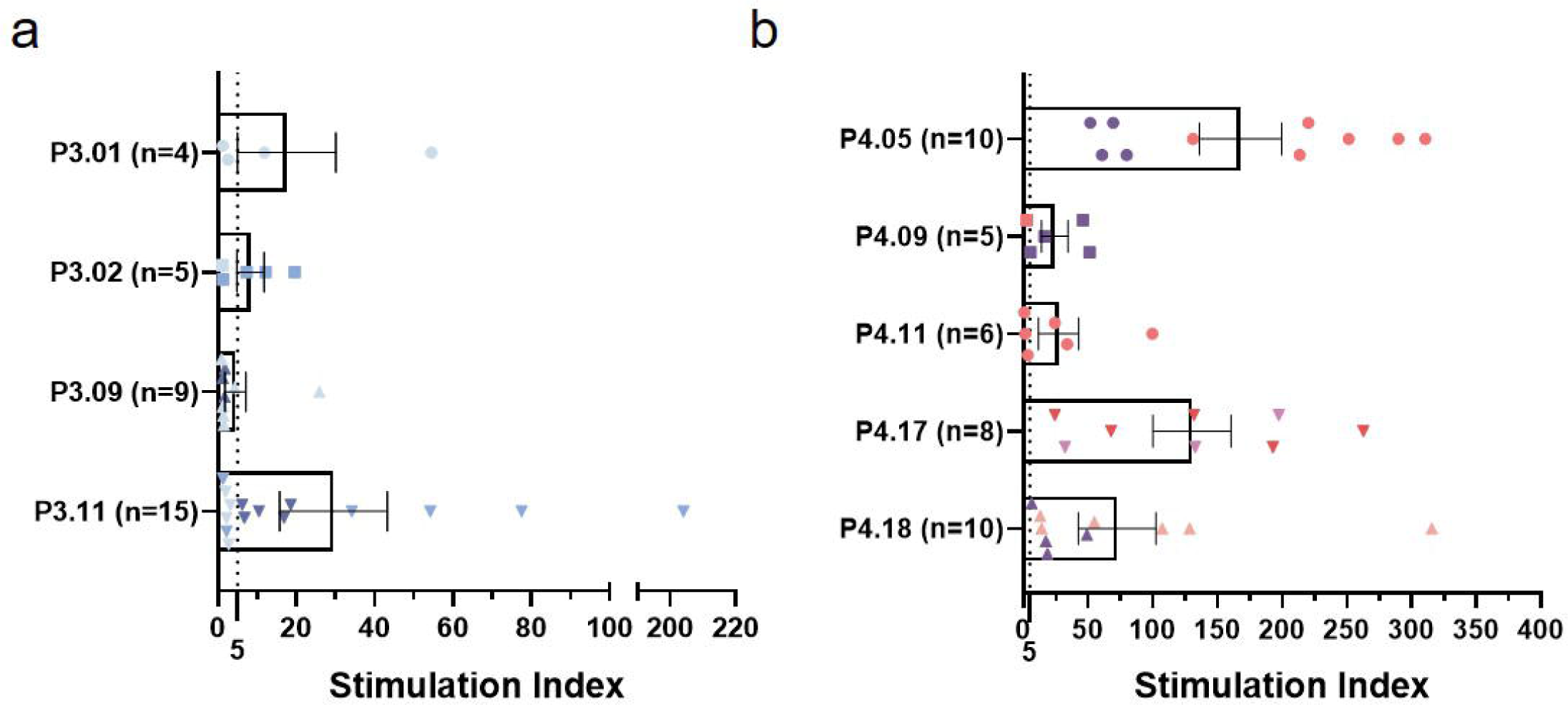
Autoreactive CD4^+^ T cell lines that recognize lncRNA-peptide epitopes are HLA-DRB1*03:01 and DRB1*04:01 restricted. Proliferation assay results represented as stimulation indexes for DRB1*03:01 peptides (a) and DRB1*04:01 peptides (b) Individual values indicate the SI for a single T cell line. The color indicates a unique original donor. SI>5, arbitrary threshold for presentation restriction proof, is depicted as a dotted line. Results are represented as means±SEM values for all the expanded lines obtained against the given peptide.

### Peptides encoded from lncRNAs are immunogenic

To analyze the immunogenic potential of high interest lncRNA-derived peptides, we performed *in silico* prediction to identify likely HLA DRB1*03:01 and HLA DRB1*04:01 binding epitopes as previously described [37]. Twelve and nineteen epitopes were predicted as potential HLA-DRB1*03:01 and HLA-DRB1*04:01 binders, respectively (**Table 2**). Most lncRNA-encoded peptides had multiple predicted epitopes and some of predicted epitopes were similar for HLA-DRB1*03:01 and HLA-DRB1*04:01 (partial overlaps between P3.05 and P4.11, and P3.09 and P4.05). An *in vitro* competitive binding assay confirmed predicted epitopes with detectable binding to HLA-DRB1*03:01 and DRB1*04:01 protein (**ESM Fig. 7a-b)**.

To assess the ability of these peptides to elicit CD4^+^ T cell responses, PBMCs from individuals with type 1 diabetes (12 HLA-DRB1*03:01 positive and 7 HLA-DRB1*04:01 positive, **ESM Table 3**) were stimulated with predicted lncRNA-peptide epitopes. After expansion, peptide reactive CD4^+^ T cells were visualized using an activation induced marker assay. Overall, ten HLA-DRB1*03:01 peptides (P3.01, P3.02, P3.04, P3.05, P3.06, P3.08, P3.09, P3.10, P3.11 and P3.12) elicited T cell expansion in at least one subject (**ESM Fig. 8**), with 7 eliciting responses in multiple donors (**Table 3**). Among these, P3.02, P3.04 and P3.09 had the highest frequency of response (**Table 3**). Overall, eight HLA-DRB1*04:01 peptides elicited T cell expansion (P4.03, P4.05, P4.07, P4.09, P4.11, P4.16, P4.17 and P4.18) (**ESM Fig. 9**), with four eliciting responses in multiple donors: P4.05, P4.09, P4.17 and P4.18 (**Table 3**).

### lncRNA peptide epitopes are presented by risk-associated HLA molecules

To confirm the HLA restriction of lncRNA-peptide epitopes, we isolated T cell lines by sorting activated cells from peptide-stimulated CD154^+^ CD69^+^ populations. We generated T cell lines for six HLA-DRB1*03:01 peptides (P3.01, P3.02, P3.05, P3.09, P3.11 and P3.12) and six HLA-DRB1*04:01 peptides (P4.05, P4.09, P4.11, P4.16, P4.17 and P4.18). Activation assays using these lines confirmed strong reactivity against the target epitope following expansion (**Fig. 3 and 4)**. We characterized the HLA restriction of the lncRNA-peptide reactive CD4^+^ T cell lines using a proliferation assay, employing single HLA molecule-expressing B cells as APCs. Restriction was assigned by comparing the SI for stimulation with peptide vs DMSO as a negative control. T cell lines recognizing P3.01, P3.02, P3.09 and P3.11 were HLA-DRB1*03:01 restricted (**Fig. 5a, ESM Table 4)**. HLA-DRB1*03:01 restricted lines recognizing P3.11 were generated from two different donors (**ESM Table 4**). DRB1*04:01 restricted lines were isolated for P4.05, P4.09, P4.11, P4.17 and P4.18 (**Fig. 5b, ESM Table 5).** Lines reactive to P4.05, P4.17, and P4.18 were isolated from multiple type 1 diabetes donors (**ESM Table 5)**.

Interestingly, donor 12, from which P3.05 T cell lines were generated (**ESM Table 3**) is heterozygous for HLA-DRB1*03:01 and HLA-DRB1*04:01. Initial epitope predictions within our candidate sequences revealed overlap between P3.05 and P4.11. Therefore, we tested those lines for both HLA-DRB1*03:01 and HLA-DRB1*04:01 restriction. Higher DRB1*04:01 SI suggested that the P3.01 T cell lines from donor 12 were, indeed, HLA-DRB1*04:01-recognized (**ESM Fig. 10a, ESM Table 4**). Conversely, in P4.11 specific lines isolated from donor 18 (HLA-DRB1*04:01 positive and HLA-DRB1*03:01 negative donor), proliferation assays with DRB1*03:01-restricted B cells showed similar SIs as DRB1*04:01-restricted B cells (**ESM Fig. 10b, ESM Table 5**), suggesting recognition of P4.11 is more likely based on the peptide sequence and TCR than the presenting HLA-DRB1 molecule.

Similarly, P3.09, partially overlapping P4.05, could give rise to both DRB1*03:01-restricted lines (donor 10) and DRB1*04:01-restricted T cell lines (donor 12) (**ESM Tables 4**).

These peptides presented by type 1 diabetes risk-associated HLA molecules constitute the most promising peptide to further characterize as diabetes relevant epitopes arising from lncRNA-peptide epitopes.

## DISCUSSION

Accumulating evidence indicates that beta cells are not passive victims of autoimmune assault. Instead, they actively contribute to their demise in through islet specific autoantigen presentation by high risk HLA molecules [5, 46]. Significant progress has been made in identifying antigens and epitopes relevant to type 1 diabetes. Neoepitopes such as hybrid insulin peptides, deamidated epitopes, and citrullinated epitopes [47–49] have been proven to be key contributors to the disease. However, the full spectrum of disease-driving epitopes remains incomplete [13, 14].

Within this context, non-coding genomic elements might be important contributors to the loss of tolerance. Several studies note that a remarkable fraction of lncRNAs associate with ribosomes [18], which is significant considering that lncRNAs are shorter and expressed at lower levels than protein coding RNAs, which may hinder the identification of ribosome associations. Patterns of ribosome protected fragments suggest that lncRNAs can translate short peptides [18, 19]. In fact, micropeptides encoded from short open reading frames within lncRNAs have been reported. Myoregulin (MLN), a 46 aa transmembrane micropeptide that regulates muscle function by inhibiting the sarcoplasmic reticulum Ca²^⁺^-ATPase (SERCA) was identified from a former lncRNA [50]. Similarly, a previously annotated lncRNA transcript, now reclassified as coding, encodes a 34 amino acid micropeptide named DWORF, which enhances SERCA activity in the cardiac muscle [51]. Most reported micropeptides have been studied in cancer, but a recent study identified a lncRNA encoded microprotein with relevance in pancreatic beta cells. The lncRNA *TUNAR* encodes a conserved 48-amino-acid micropeptide named BNLN (beta cell- and neural cell-regulin) which localizes to the ER in beta cells and plays a crucial role in maintaining calcium homeostasis[24].

The number of studies showcasing lncRNA translation is increasing, but no prior work has associated micropeptide translation with autoimmunity in beta cells. Viral and proinflammatory insults induce profound changes in lncRNA transcriptome in beta cells[19, 39], provoking upregulation of many transcripts. This observation is replicated in our ribosome associated RNAseq data and aligns with previous data[19], suggesting an increase of ribosome-bound lncRNA, potentially leading to non-canonical translation events. In parallel, HLA expression is upregulated in response to inflammatory stimuli in pancreatic beta cells[52–54], expanding the epitope landscape and arguably promoting the presentation of epitopes and neoepitopes, such as lncRNA derived peptides, that likely circumvent thymic tolerance mechanisms[55].

We have combined ribosome associated RNA sequencing with mass spectrometry to identify lncRNA candidates with coding potential. Although our ribosome-bound RNA sequencing strategy is less resource intensive than standard ribosome profiling, it lacks the ribosome occupancy information, which is helpful to confirm that lncRNA transcripts with ribosomal footprints are actively being translated rather than spurious or regulatory ribosome-lncRNA associations. With respect to the nature of the start codon, translation outside of annotated protein-coding regions is relatively rare and often independent of classical AUG start codons, but emerging evidence highlights the role of near-cognate codons (such as CUG, GUG, etc.) in initiating translation of short open reading frames [56]. Nevertheless, we focused on canonical AUG-initiated ORFs and therefore excluded near-cognate initiated ORFs from our custom translation database, allowing for a more targeted analysis while leaving room for future exploration of alternative initiation mechanisms.

In our study, *UXT-AS1*- and *RAPGEF4-AS1*-encoded peptides were detected in pancreatic beta cells under basal conditions; the peptide encoded by lncRNA *ENSG00000227066* was only observed in beta cells treated with PIC. These findings suggest that some peptides are produced constitutively in pancreatic beta cells, whereas others are induced by inflammation. Future work should elucidate the regulatory mechanisms controlling translation of these lncRNAs. Our results suggest that features such as IRES and m^6^A modifications may not be absolutely required for translation; only *ENSG00000227066* was m^6^A-methylated in PIC-transfected pancreatic beta cells. However, m^6^A-methylation may increase abundance, given that only this lncRNA was detected in the MS analysis of PIC-transfected beta cells.

Multiple proteomic studies of beta cells have been performed, but peptides translated from lncRNA have not been previously reported. Their absence from previous proteomic datasets is likely due to a combination of methodological and biological factors [57–59]. Although two of the three peptides (90–214 amino acids) exceed the standard 100 amino acid cutoff for micropeptide designation, they are still relatively small and likely expressed at low abundance, which reduces their detectability in conventional workflows. The use of active nascent peptide mass spectrometry in our study enabled the enrichment and detection of low-abundance, newly synthesized proteins, revealing translated peptides that were previously undetected.

In this work we have shown evidence that lncRNAs can encode peptides and that circulating CD4^+^ T cells from can recognize them as epitopes. To our knowledge, these could be the first type 1 diabetes neoepitopes described that are generated through the translation of non-coding elements of the genome. The discovery that lncRNA-derived peptides can be produced and translated by beta cells and recognized by T cells expands the conventional view of the beta cell immunopeptidome. The translational capacity of lncRNAs, once thought to be exclusively regulatory, is increasingly recognized across tissues and disease contexts. For example, a proteogenomic study of two murine cancer cell lines and seven human primary tumors revealed that around 90% of tumor-specific antigens are derived from non-coding regions [60]. Similar results have been found in hepatocellular carcinoma, in which 33 tumor-specific lncRNAs encoding novel cancer antigens were identified [61].

In summary, in this study we have identified a novel class of autoantigens derived from peptides encoded by three lncRNAs (*UXT-AS1*, *RAPGEF4-AS1* and *ENSG00000227066)* that are translated in human pancreatic beta cells and can elicit autoreactive CD4^⁺^ T cell responses in individuals with type 1 diabetes. Our findings provide direct evidence that lncRNA-derived peptides can act as a previously unrecognized source of autoantigens, broadening the landscape of potential neoepitopes contributing to beta cell–specific immune responses in type 1 diabetes. Further studies to characterize the frequency and phenotype of such autoreactive T cells in healthy controls versus individuals with type 1 diabetes are needed to clarify their relevance and evaluate their potential use as a biomarker for disease progression or disease subtype stratification.

## ACKNOWLEDGEMENTS

The authors want to thank Hai Nguyen (Benaroya Research Institute) for his insights in successful PBMC cultures. We would also like to thank to the SGIKER UPV-EHU services for their technical support and expertise in proteomics and microscopy experiments.

## DATA AVAILABILITY

RNA-seq data are available in Sequence Read Archive (SRA) under accession number PRJNA1366636 and mass spectrometry data are available upon request to the authors.

## FUNDING

This work was supported by the Spanish Ministry of Science, Innovation and Universities (PID2023-150479NB-I00) and the Basque Department of Health (2021111001) to IS. JMS and LBM are supported by Predoctoral Fellowship Grants from the Education department of the Basque Government. IP is supported by a Predoctoral Fellowship Grant from Spanish Ministry of Science, Innovation and Universities.

## AUTHOR’S RELATIONSHIPS AND ACTIVITIES

No potential conflicts of interest relevant to this article were reported.

## CONTRIBUTION STATEMENT

JMS contributed to research data, wrote, revised and edited the manuscript. AC, KGE, LB, IP, ARJ, KTL and HRM contributed to researched data, and revised and edited the manuscript. ACR and EAJ contributed to the design and interpretation of the experiments, contributed to discussion, and revised and edited the manuscript. IS contributed to the original idea, design and interpretation of experiments, contributed to discussion, and wrote, revised, and edited the manuscript. All authors have read and approved the manuscript. IS is the guarantor of this work and, as such, has full access to all the data in the study and takes responsibility for the integrity of the data and the accuracy of the data analysis.

## ABBREVIATIONS

APC: Antigen presenting cell
AHA: Azidohomoalanine
DRiPs: Defective ribosomal products
ER: Endoplasmic reticulum
HLA: Human Leucocyte Antigen lnc
RNA: Long non-coding RNA
ORF: Open reading frame
PBMC: Peripheral blood mononuclear cell
PIC: Polyinosinic:polycytidylic acid
SI: Stimulation index

## Notes

### Competing Interest Statement

The authors have declared no competing interest.

## REFERENCES

1. Katsarou A, Gudbjörnsdottir S, Rawshani A, et al (2017) Type 1 diabetes mellitus. Nat Rev Dis Primers 3(1):17016. 10.1038/nrdp.2017.16

2. Ziegler AG, Rewers M, Simell O, et al (2013) Seroconversion to multiple islet autoantibodies and risk of progression to diabetes in children. JAMA 309(23):2473–9. 10.1001/jama.2013.6285

3. Krischer JP, Lynch KF, Schatz DA, et al (2015) The 6 year incidence of diabetes-associated autoantibodies in genetically at-risk children: the TEDDY study. Diabetologia 58(5):980–987. 10.1007/s00125-015-3514-y

4. Marré ML, James EA, Piganelli JD (2015) β cell ER stress and the implications for immunogenicity in type 1 diabetes. Front Cell Dev Biol 3. 10.3389/fcell.2015.00067

5. Mallone R, Eizirik DL (2020) Presumption of innocence for beta cells: why are they vulnerable autoimmune targets in type 1 diabetes? Diabetologia 63(10):1999–2006. 10.1007/s00125-020-05176-7

6. Callebaut A, Guyer P, Derua R, et al (2024) CD4+ T Cells From Individuals With Type 1 Diabetes Respond to a Novel Class of Deamidated Peptides Formed in Pancreatic Islets. Diabetes 73(5):728–742. 10.2337/db23-0588

7. Bender C, Rodriguez-Calvo T, Amirian N, Coppieters KT, von Herrath MG (2020) The healthy exocrine pancreas contains preproinsulin-specific CD8 T cells that attack islets in type 1 diabetes. Sci Adv 6(42). 10.1126/sciadv.abc5586

8. Rodriguez-Calvo T, Zapardiel-Gonzalo J, Amirian N, et al (2017) Increase in Pancreatic Proinsulin and Preservation of β-Cell Mass in Autoantibody-Positive Donors Prior to Type 1 Diabetes Onset. Diabetes 66(5):1334–1345. 10.2337/db16-1343

9. Noble JA, Valdes AM, Varney MD, et al (2010) HLA Class I and Genetic Susceptibility to Type 1 Diabetes. Diabetes 59(11):2972–2979. 10.2337/db10-0699

10. Vita R, Blazeska N, Marrama D, et al (2025) The Immune Epitope Database (IEDB): 2024 update. Nucleic Acids Res 53(D1):D436–D443. 10.1093/nar/gkae1092

11. James EA, Mallone R, Kent SC, DiLorenzo TP (2020) T-Cell Epitopes and Neo-epitopes in Type 1 Diabetes: A Comprehensive Update and Reappraisal. Diabetes 69(7):1311–1335. 10.2337/dbi19-0022

12. Rodriguez-Calvo T, Johnson JD, Overbergh L, Dunne JL (2021) Neoepitopes in Type 1 Diabetes: Etiological Insights, Biomarkers and Therapeutic Targets. Front Immunol 12:667989. 10.3389/fimmu.2021.667989

13. Piganelli JD, Mamula MJ, James EA (2021) The Role of β Cell Stress and Neo-Epitopes in the Immunopathology of Type 1 Diabetes. Front Endocrinol (Lausanne) 11. 10.3389/fendo.2020.624590

14. James EA, Mallone R, Kent SC, DiLorenzo TP (2020) T-Cell Epitopes and Neo-epitopes in Type 1 Diabetes: A Comprehensive Update and Reappraisal. Diabetes 69(7):1311–1335. 10.2337/dbi19-0022

15. James EA, Pietropaolo M, Mamula MJ (2018) Immune Recognition of β-Cells: Neoepitopes as Key Players in the Loss of Tolerance. Diabetes 67(6):1035–1042. 10.2337/dbi17-0030

16. Coppieters KT, Dotta F, Amirian N, et al (2012) Demonstration of islet-autoreactive CD8 T cells in insulitic lesions from recent onset and long-term type 1 diabetes patients. Journal of Experimental Medicine 209(1):51–60. 10.1084/jem.20111187

17. Zeng C, Fukunaga T, Hamada M (2018) Identification and analysis of ribosome-associated lncRNAs using ribosome profiling data. BMC Genomics 19(1). 10.1186/s12864-018-4765-z

18. Ingolia NT, Brar GA, Stern-Ginossar N, et al (2014) Ribosome Profiling Reveals Pervasive Translation Outside of Annotated Protein-Coding Genes. Cell Rep 8(5):1365–1379. 10.1016/j.celrep.2014.07.045

19. Thomaidou S, Slieker RC, van der Slik AR, et al (2021) Long RNA Sequencing and Ribosome Profiling of Inflamed β-Cells Reveal an Extensive Translatome Landscape. Diabetes 70(10):2299–2312. 10.2337/db20-1122

20. Ylipaasto P, Kutlu B, Rasilainen S, et al (2005) Global profiling of coxsackievirus-and cytokine-induced gene expression in human pancreatic islets. Diabetologia 48(8):1510–1522. 10.1007/s00125-005-1839-7

21. Statello L, Guo CJ, Chen LL, Huarte M (2021) Gene regulation by long non-coding RNAs and its biological functions. Nat Rev Mol Cell Biol 22:96–118

22. Ruiz-Orera J, Messeguer X, Subirana JA, Alba MM (2014) Long non-coding RNAs as a source of new peptides. Elife 3. 10.7554/eLife.03523

23. Senís E, Esgleas M, Najas S, et al (2021) TUNAR lncRNA Encodes a Microprotein that Regulates Neural Differentiation and Neurite Formation by Modulating Calcium Dynamics. Front Cell Dev Biol 9. 10.3389/fcell.2021.747667

24. Li M, Shao F, Qian Q, et al (2021) A putative long noncoding RNA-encoded micropeptide maintains cellular homeostasis in pancreatic β cells. Mol Ther Nucleic Acids 26:307–320. 10.1016/j.omtn.2021.06.027

25. Bolger AM, Lohse M, Usadel B (2014) Trimmomatic: a flexible trimmer for Illumina sequence data. Bioinformatics 30(15):2114–2120. 10.1093/bioinformatics/btu170

26. Kim D, Paggi JM, Park C, Bennett C, Salzberg SL (2019) Graph-based genome alignment and genotyping with HISAT2 and HISAT-genotype. Nat Biotechnol 37(8):907–915. 10.1038/s41587-019-0201-4

27. Pertea M, Pertea GM, Antonescu CM, Chang T-C, Mendell JT, Salzberg SL (2015) StringTie enables improved reconstruction of a transcriptome from RNA-seq reads. Nat Biotechnol 33(3):290–295. 10.1038/nbt.3122

28. Anders S, Pyl PT, Huber W (2015) HTSeq—a Python framework to work with high-throughput sequencing data. Bioinformatics 31(2):166–169. 10.1093/bioinformatics/btu638

29. Robinson MD, McCarthy DJ, Smyth GK (2010) *edgeR*□: a Bioconductor package for differential expression analysis of digital gene expression data. Bioinformatics 26(1):139–140. 10.1093/bioinformatics/btp616

30. Chen EY, Tan CM, Kou Y, et al (2013) Enrichr: interactive and collaborative HTML5 gene list enrichment analysis tool. BMC Bioinformatics 14(1):128. 10.1186/1471-2105-14-128

31. Minati L, Firrito C, Del Piano A, et al (2021) One-shot analysis of translated mammalian lncRNAs with AHARIBO. Elife 10. 10.7554/eLife.59303

32. Castellanos-Rubio A, Fernandez-Jimenez N, Kratchmarov R, et al (2016) A long noncoding RNA associated with susceptibility to celiac disease. Science (1979) 352(6281):91–95. 10.1126/science.aad0467

33. Fan R, Cui C, Kang B, Chang Z, Wang G, Cui Q (2024) A combined deep learning framework for mammalian m6A site prediction. Cell Genomics 4(12):100697. 10.1016/j.xgen.2024.100697

34. Jumper J, Evans R, Pritzel A, et al (2021) Highly accurate protein structure prediction with AlphaFold. Nature 596(7873):583–589. 10.1038/s41586-021-03819-2

35. Walker JM (2005) The Proteomics Protocols Handbook. Humana Press, Totowa, NJ

36. Thumuluri V, Almagro Armenteros JJ, Johansen AR, Nielsen H, Winther O (2022) DeepLoc 2.0: multi-label subcellular localization prediction using protein language models. Nucleic Acids Res 50(W1):W228–W234. 10.1093/nar/gkac278

37. Ettinger RA, Papadopoulos GK, Moustakas AK, Nepom GT, Kwok WW (2006) Allelic Variation in Key Peptide-Binding Pockets Discriminates between Closely Related Diabetes-Protective and Diabetes-Susceptible *HLA-DQB1*06* Alleles. The Journal of Immunology 176(3):1988–1998. 10.4049/jimmunol.176.3.1988

38. Kovats S, Nepom GT, Coleman M, Nepom B, Kwok WW, Blum JS (1995) Deficient antigen-presenting cell function in multiple genetic complementation groups of type II bare lymphocyte syndrome. Journal of Clinical Investigation 96(1):217–223. 10.1172/JCI118023

39. Santin I, Eizirik DL (2013) Candidate genes for type 1 diabetes modulate pancreatic islet inflammation and β -cell apoptosis. Diabetes Obes Metab 15(s3):71–81. 10.1111/dom.12162

40. Szafron LM, Balcerak A, Grzybowska EA, et al (2015) The Novel Gene CRNDE Encodes a Nuclear Peptide (CRNDEP) Which Is Overexpressed in Highly Proliferating Tissues. PLoS One 10(5):e0127475. 10.1371/journal.pone.0127475

41. Zhang M, Zhao K, Xu X, et al (2018) A peptide encoded by circular form of LINC-PINT suppresses oncogenic transcriptional elongation in glioblastoma. Nat Commun 9(1):4475. 10.1038/s41467-018-06862-2

42. Dang HX, Saha D, Jayasinghe R, et al (2023) Single-cell transcriptomics reveals long noncoding RNAs associated with tumor biology and the microenvironment in pancreatic cancer. NAR Cancer 5(4). 10.1093/narcan/zcad055

43. Uddin MB, Wang Z, Yang C (2025) Epitranscriptomic RNA m ^6^ A Modification in Cancer Therapy Resistance: Challenges and Unrealized Opportunities. Advanced Science 12(4). 10.1002/advs.202403936

44. Zhang X-N, Yang K-D, Chen C, et al (2021) Pericytes augment glioblastoma cell resistance to temozolomide through CCL5-CCR5 paracrine signaling. Cell Res 31(10):1072–1087. 10.1038/s41422-021-00528-3

45. Fan R, Cui C, Kang B, Chang Z, Wang G, Cui Q (2024) A combined deep learning framework for mammalian m6A site prediction. Cell Genomics 4(12):100697. 10.1016/j.xgen.2024.100697

46. Roep BO, Thomaidou S, van Tienhoven R, Zaldumbide A (2021) Type 1 diabetes mellitus as a disease of the β-cell (do not blame the immune system?). Nat Rev Endocrinol 17(3):150–161. 10.1038/s41574-020-00443-4

47. McGinty JW, Marré ML, Bajzik V, Piganelli JD, James EA (2015) T Cell Epitopes and Post-Translationally Modified Epitopes in Type 1 Diabetes. Curr Diab Rep 15(11):90. 10.1007/s11892-015-0657-7

48. Crawford SA, Wiles TA, Wenzlau JM, et al (2022) Cathepsin D Drives the Formation of Hybrid Insulin Peptides Relevant to the Pathogenesis of Type 1 Diabetes. Diabetes 71(12):2793–2803. 10.2337/db22-0303

49. Azoury ME, Tarayrah M, Afonso G, et al (2020) Peptides Derived From Insulin Granule Proteins Are Targeted by CD8+ T Cells Across MHC Class I Restrictions in Humans and NOD Mice. Diabetes 69(12):2678–2690. 10.2337/db20-0013

50. Anderson DM, Anderson KM, Chang C-L, et al (2015) A Micropeptide Encoded by a Putative Long Noncoding RNA Regulates Muscle Performance. Cell 160(4):595–606. 10.1016/j.cell.2015.01.009

51. Nelson BR, Makarewich CA, Anderson DM, et al (2016) A peptide encoded by a transcript annotated as long noncoding RNA enhances SERCA activity in muscle. Science (1979) 351(6270):271–275. 10.1126/science.aad4076

52. Russell MA, Redick SD, Blodgett DM, et al (2019) HLA Class II Antigen Processing and Presentation Pathway Components Demonstrated by Transcriptome and Protein Analyses of Islet β-Cells From Donors With Type 1 Diabetes. Diabetes 68(5):988–1001. 10.2337/db18-0686

53. Quesada-Masachs E, Zilberman S, Rajendran S, et al (2022) Upregulation of HLA class II in pancreatic beta cells from organ donors with type 1 diabetes. Diabetologia 65(2):387–401. 10.1007/s00125-021-05619-9

54. Richardson SJ, Rodriguez-Calvo T, Gerling IC, et al (2016) Islet cell hyperexpression of HLA class I antigens: a defining feature in type 1 diabetes. Diabetologia 59(11):2448–2458. 10.1007/s00125-016-4067-4

55. Nanaware PP, Calvo-Calle JM, Redick SD, et al (2025) The antigen presentation landscape of cytokine-stressed human pancreatic islets. Cell Rep 44(8):115927. 10.1016/j.celrep.2025.115927

56. Andreev DE, Loughran G, Fedorova AD, Mikhaylova MS, Shatsky IN, Baranov P V. (2022) Non-AUG translation initiation in mammals. Genome Biol 23(1):111. 10.1186/s13059-022-02674-2

57. Woo J, Sudhir P, Zhang Q (2020) Pancreatic Tissue Proteomics Unveils Key Proteins, Pathways, and Networks Associated with Type 1 Diabetes. Proteomics Clin Appl 14(6). 10.1002/prca.202000053

58. Lee J, Wu Y, Schnepp P, et al (2015) Proteomics analysis of rough endoplasmic reticulum in pancreatic beta cells. Proteomics 15(9):1508–1511. 10.1002/pmic.201400345

59. Metz TO, Jacobs JM, Gritsenko MA, et al (2006) Characterization of the Human Pancreatic Islet Proteome by Two-Dimensional LC/MS/MS. J Proteome Res 5(12):3345–3354. 10.1021/pr060322n

60. Laumont CM, Vincent K, Hesnard L, et al (2018) Noncoding regions are the main source of targetable tumor-specific antigens. Sci Transl Med 10(470). 10.1126/scitranslmed.aau5516

61. Camarena ME, Theunissen P, Ruiz M, et al (2024) Microproteins encoded by noncanonical ORFs are a major source of tumor-specific antigens in a liver cancer patient meta-cohort. Sci Adv 10(28). 10.1126/sciadv.adn3628

