## Supplementary material for "Long non-coding RNAs as a novel source of beta cell autoantigens in type 1 diabetes"

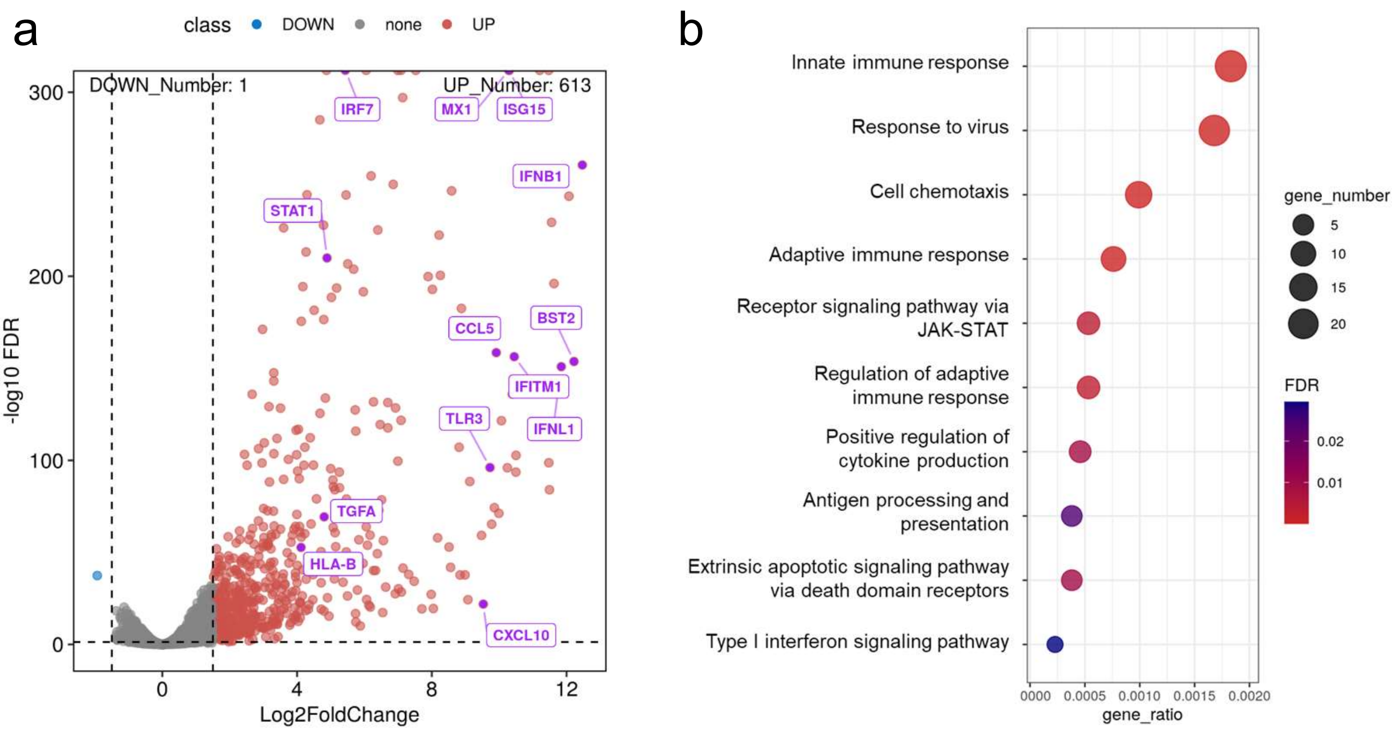


**ESM Fig. 1. Intracellular PIC induced upregulation of antiviral response and proinflammatory pathway genes in pancreatic beta cells**. A) Volcano plot of the differentially expressed protein coding genes upon intracellular PIC exposure. B) KEGG pathway analysis of the significantly upregulated protein coding gene subset.

a

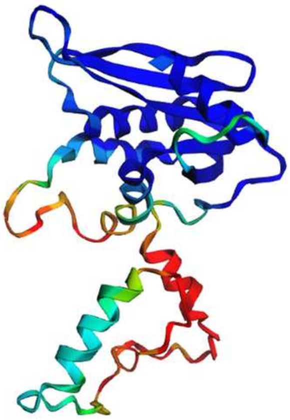

**Sequence**
**Predicted localizations:** Cytoplasm, Nucleus
**Predicted signals:** Nuclear localization signal, Nuclear export signal

| Localization | Cytoplasm | Nucleus | Extracellular | Cell membrane | Mitochondrion | Plastid | Endoplasmic reticulum | Lysosome/ Vacuole | Golgi apparatus | Peroxisome |
|---|---|---|---|---|---|---|---|---|---|---|
| Probability | 0.7610 | 0.8270 | 0.1832 | 0.1853 | 0.1560 | 0.0485 | 0.1925 | 0.0273 | 0.0792 | 0.0632 |

b

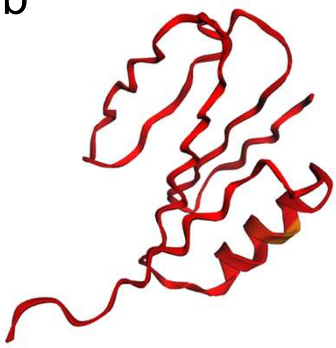

**Sequence**
**Predicted localizations:** Nucleus, Mitochondrion
**Predicted signals:**

| Localization | Cytoplasm | Nucleus | Extracellular | Cell membrane | Mitochondrion | Plastid | Endoplasmic reticulum | Lysosome/ Vacuole | Golgi apparatus | Peroxisome |
|---|---|---|---|---|---|---|---|---|---|---|
| Probability | 0.4405 | 0.5642 | 0.1281 | 0.0838 | 0.7120 | 0.0034 | 0.1312 | 0.1196 | 0.1548 | 0.2882 |

c

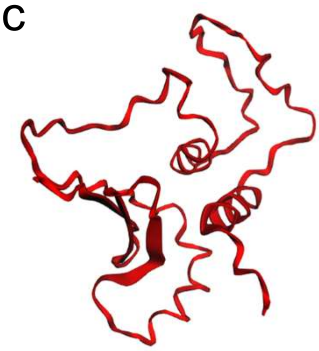

**Sequence**
**Predicted localizations:** Cytoplasm
**Predicted signals:** Nuclear localization signal

| Localization | Cytoplasm | Nucleus | Extracellular | Cell membrane | Mitochondrion | Plastid | Endoplasmic reticulum | Lysosome/ Vacuole | Golgi apparatus | Peroxisome |
|---|---|---|---|---|---|---|---|---|---|---|
| Probability | 0.6797 | 0.4894 | 0.1937 | 0.1191 | 0.5411 | 0.0046 | 0.2184 | 0.0444 | 0.1398 | 0.0041 |

**ESM Fig. 2. Predicted structural models and subcellular localization of the peptides encoded by lncRNAs *ENSG00000227066*, *UXT-AS1* and *RAPGEF4-AS1*.** AlphaFold structural predictions of the newly identified peptides encoded from *ENSG00000227066* (a) , *UXT-AS1* (b) and *RAPGEF4-AS1* (c) are shown, colored according to the per-residue confidence score (pLDDT). Dark blue regions correspond to very high confidence predictions (pLDDT>90), followed by cyan (pLDDT=80), green (pLDDT=70), yellow (pLDDT=60) and red (pLDDT<50). Subcellular localizations of the peptides predicted by DeepLoc are shown in the tables. In green are highlighted the most probable localization for each peptide.

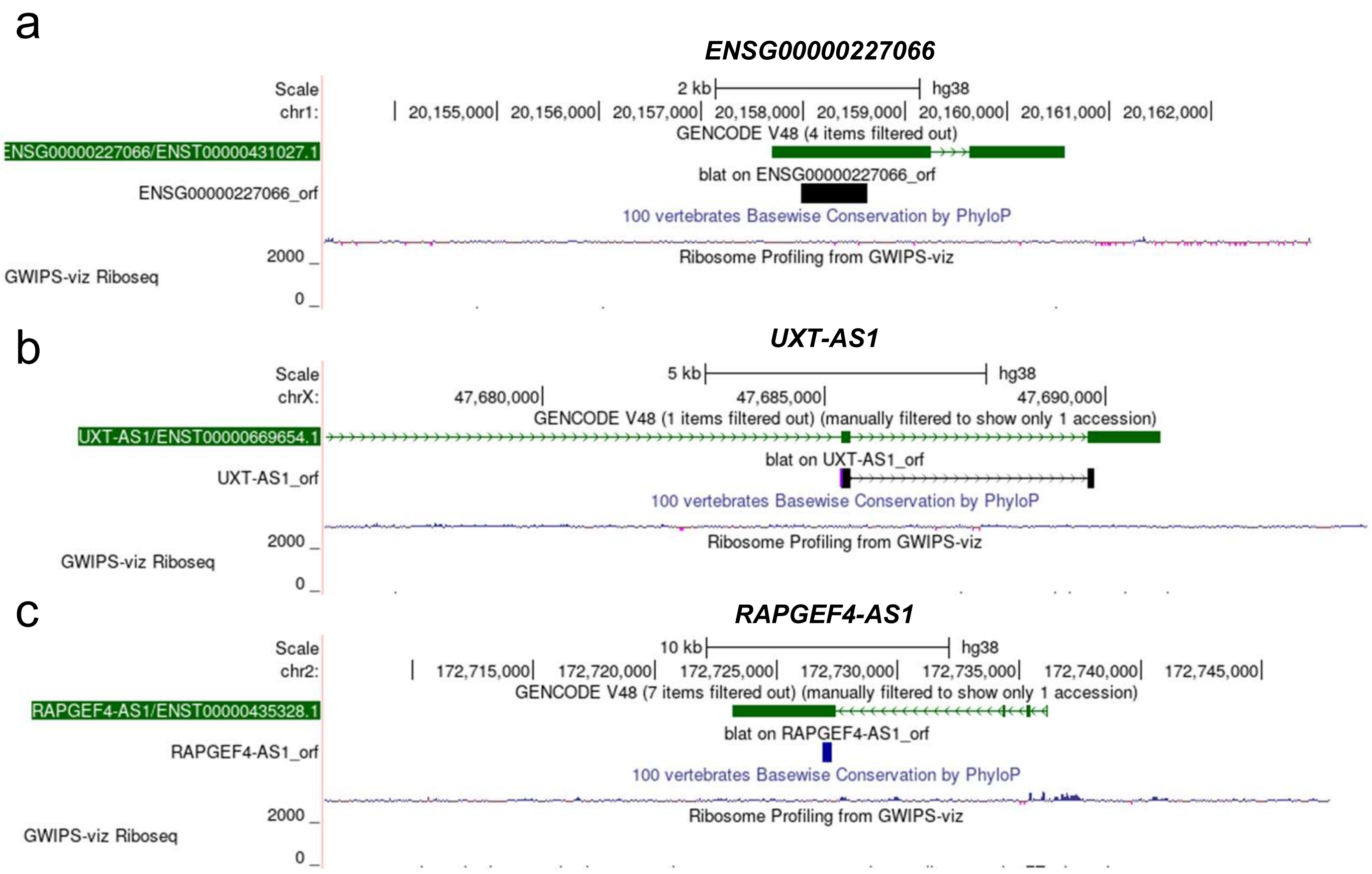


**ESM Fig. 3. Genomic location of the T1D neoantigen-encoding lncRNAs.** a-c) UCSC browser snapshots of the three lncRNA candidates. There is no noticeable base wise conservation in any of the three candidates according to PhyloP conservation track. Additionally, there is not significant coverage data of the transcripts of interest in the GWIPS Riboseq read database browser.

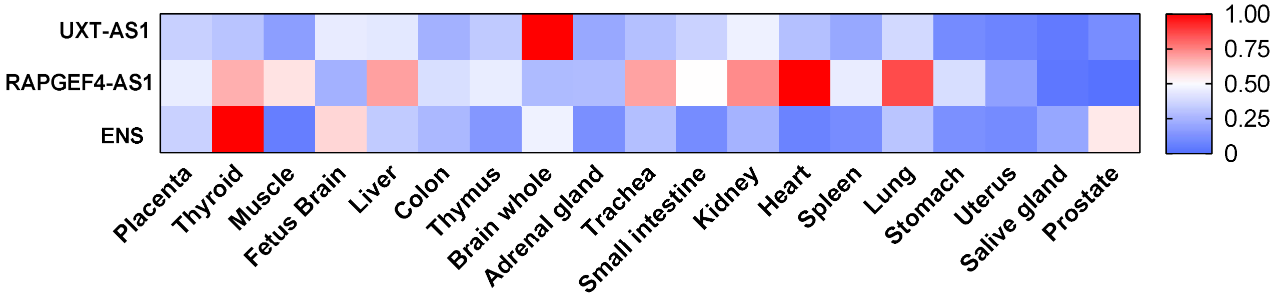


**ESM Fig. 4. Peptide-encoding lncRNAs are ubiquitously expressed in human tissues at low expression values.** Expression of *UXT-AS1* (upper panel), *RAPGEF4-AS1* (middle panel) and *ENSG00000227066* (ENS; lower panel) in a set of human tissues. The mean expression level of three experimental replicates is shown as a heat map. All expression values were normalized to the highest to get values between 0 and 1.

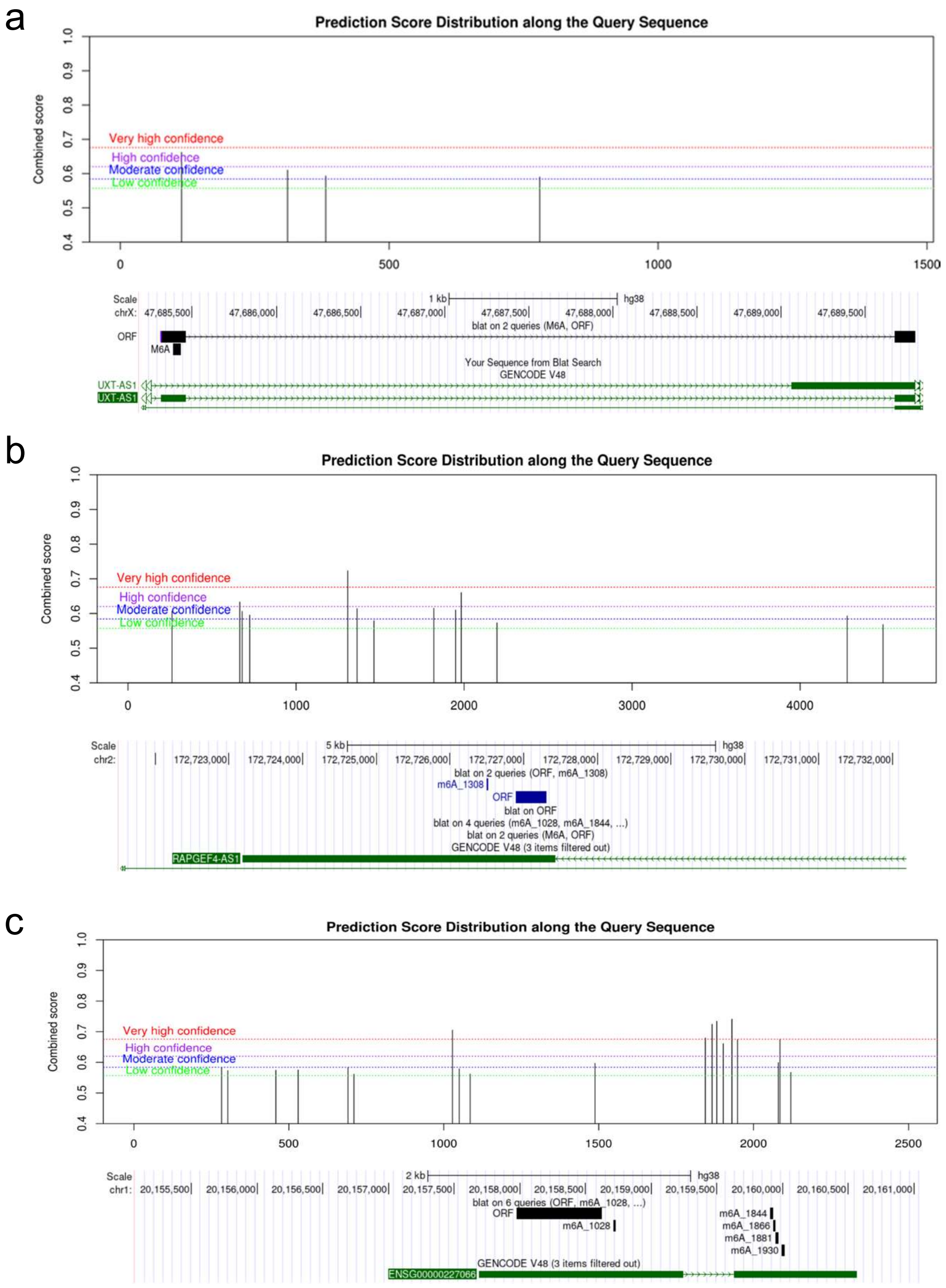


**ESM Fig. 5. m6A methylation prediction revealed several potential m6A motifs in UXT-AS1, RAPGEF-AS1 and ENSG00000227066 transcripts**. m6A methylation motifs were predicted in UXT-AS1 (a), RAPGEF-AS1 (b) and ENSG00000227066 (c) transcripts using SRAMP prediction tool. One "very high confidence" mark was predicted in UXT-AS1 (a) and RAPGEF-AS1 (b), and five "very high confidence" marks were predicted in ENSG00000227066 (c).

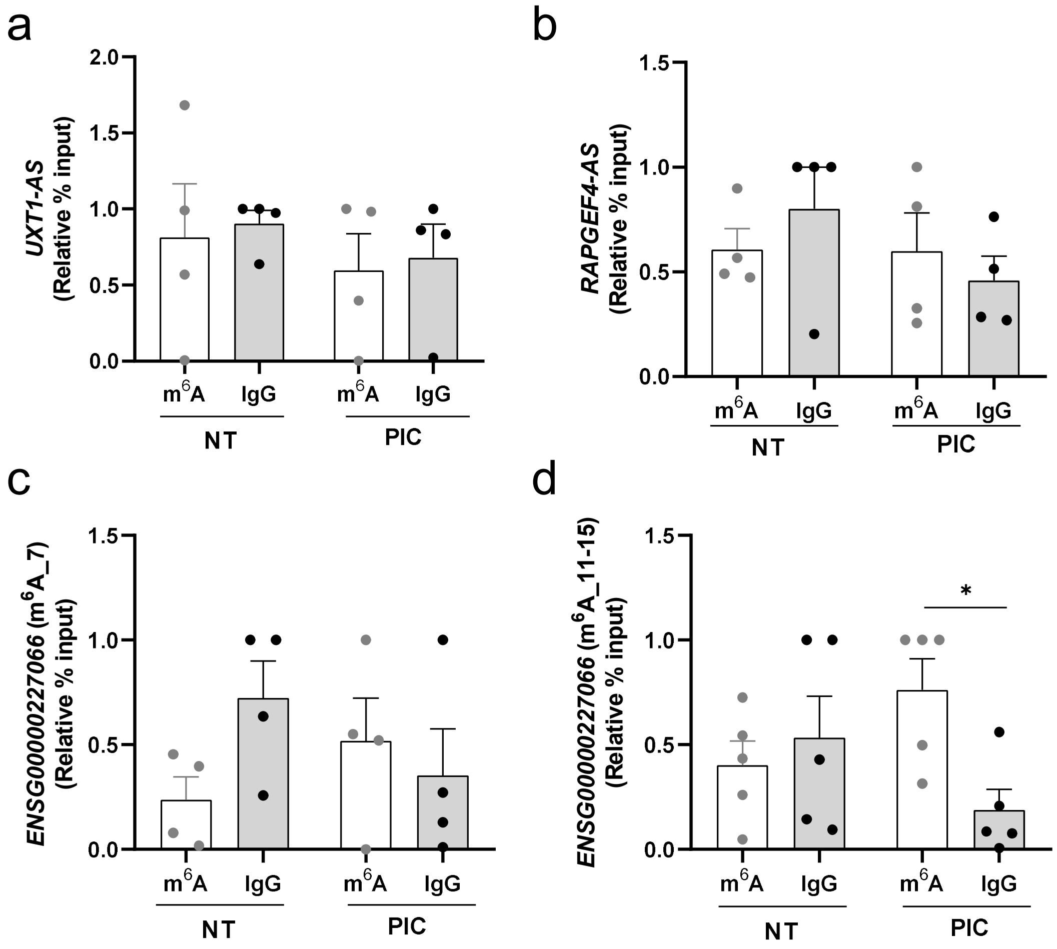


**ESM Fig. 6. LncRNA *ENSG00000227066* transcript is m6A-methylated upon intracellular PIC exposure in pancreatic beta cells**. m6A methylation was determined in basal (NT) and PIC-transfected cells (PIC) by m6A-RIP followed by a qPCR of *UXT-AS1* (a), *RAPGEF4-AS1* (b) and *ENSG00000227066* (c-d). While *UXT-AS1* (a), *RAPGEF4-AS1* (b), and the region m6A_7 of *ENSG0000022706* (c) were not methylated, m6A_11-15 region of *ENSG00000227066* transcript (d) was unmethylated at basal conditions but significantly methylated following intracellular PIC exposure. *p<0.05; Student's t test.

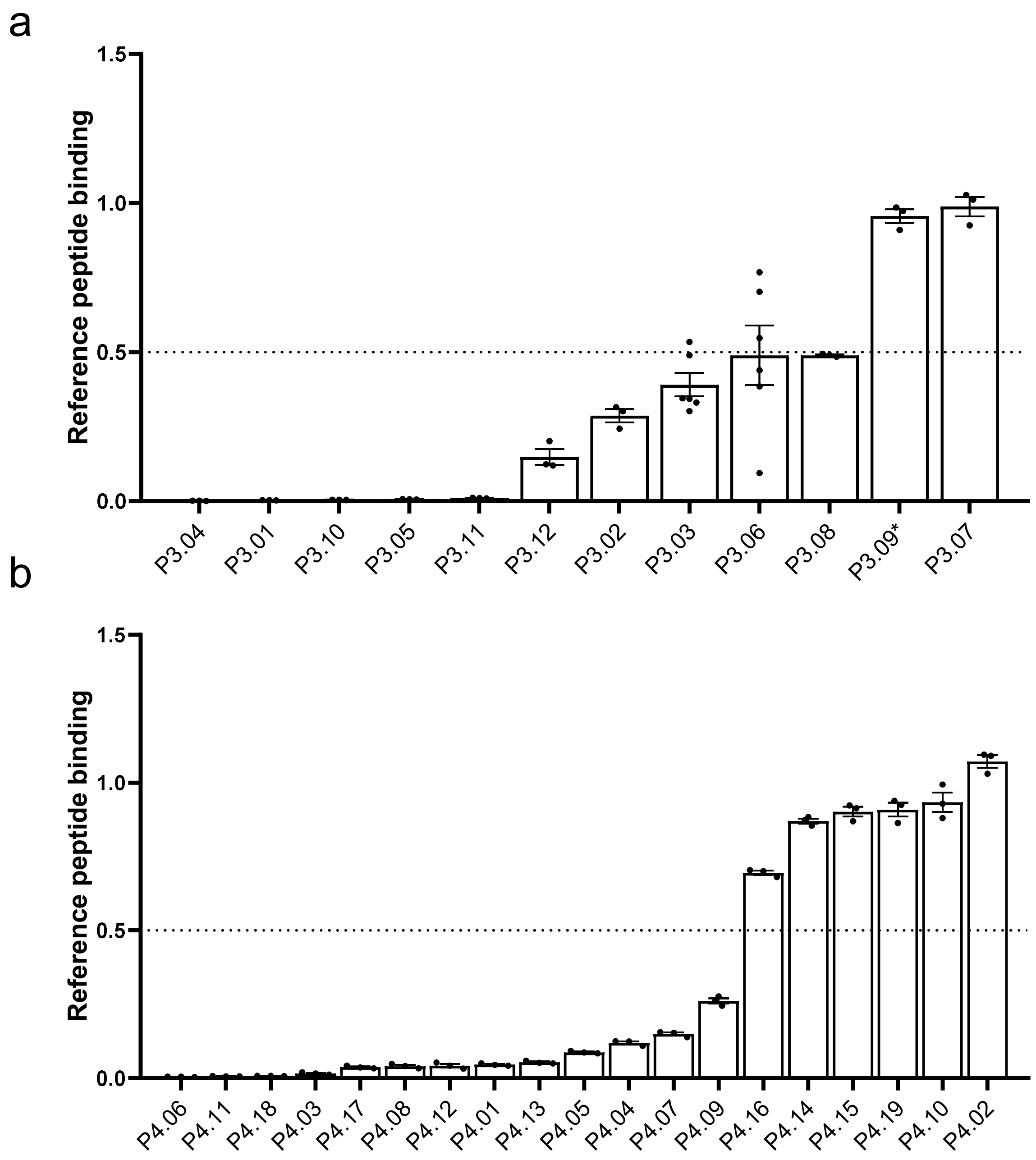


**ESM Fig. 7. Several predicted epitopes within the lncRNAs bind to HLA-DRB1*03:01 or HLA-DRB1*04:01 molecules. C**ompetitive affinity binding assay was performed in triplicate (except P3.03 and P3.06 n=6), data represented as mean±SEM, less is more. More than half of the predicted epitopes showed substantial binding to the HLA-DRB1*03:01 (a) and HLA-DRB1*04:01 (b) molecules (relative reference peptide binding <0.5). *Note that P3.09, which was not a strong binder, was selected for in vitro studies due to peptide synthesis delays, bringing the final number of selected peptides for further studies up to 11 for HLA-DRB1*03:01 and 13 for HLA-DRB1*04:01.

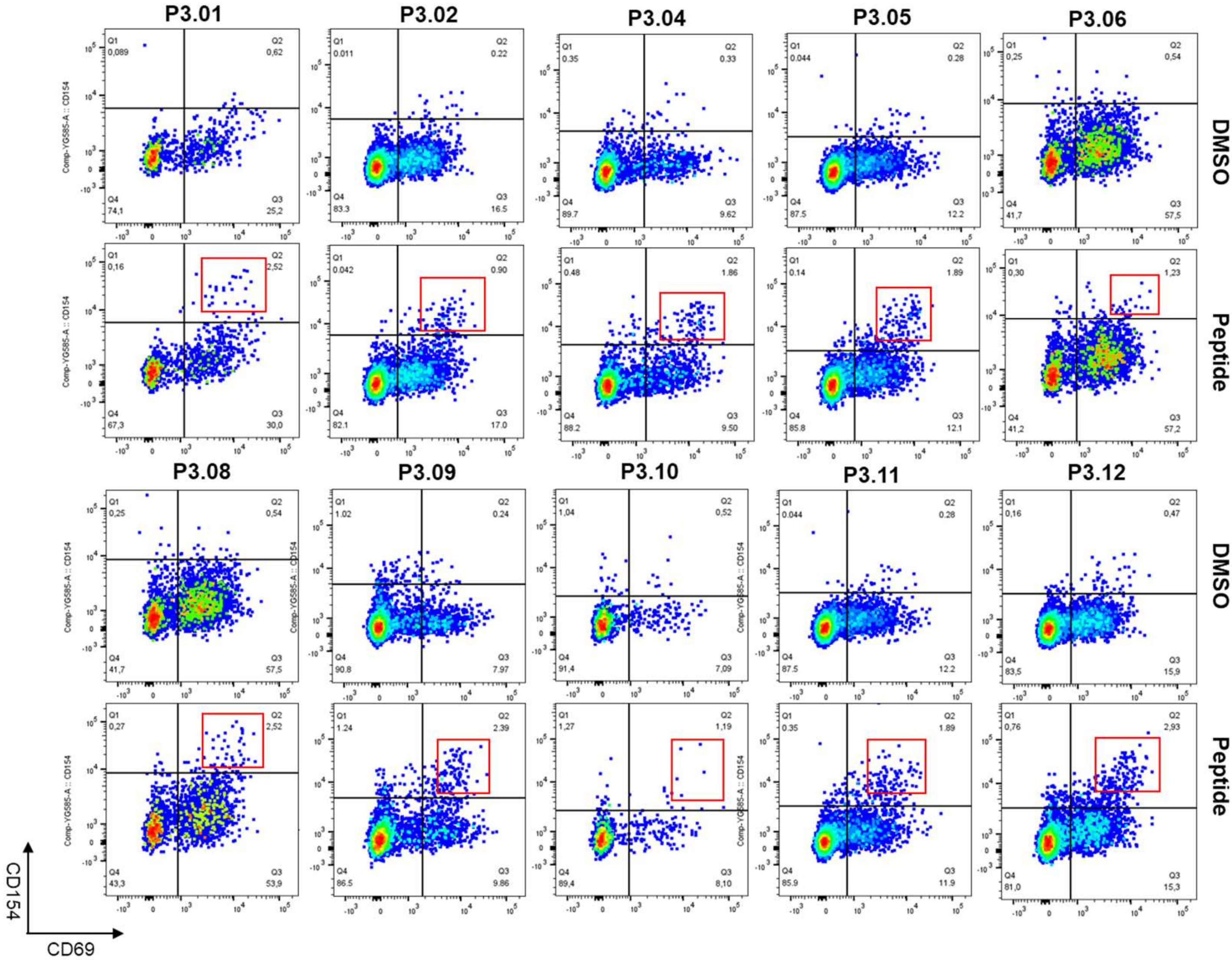


**ESM Fig. 8. Representative FACS plots of positive CD154 AIM responses for each HLA-DRB1*03:01 peptide**. Ten epitopes (P3.01, P3.02, P3.04, P3.05, P3.06, P3.08, P3.09, P3.10, P3.11 and P3.12) induced activation of T cells in at least one donor. DMSO was used as negative control.

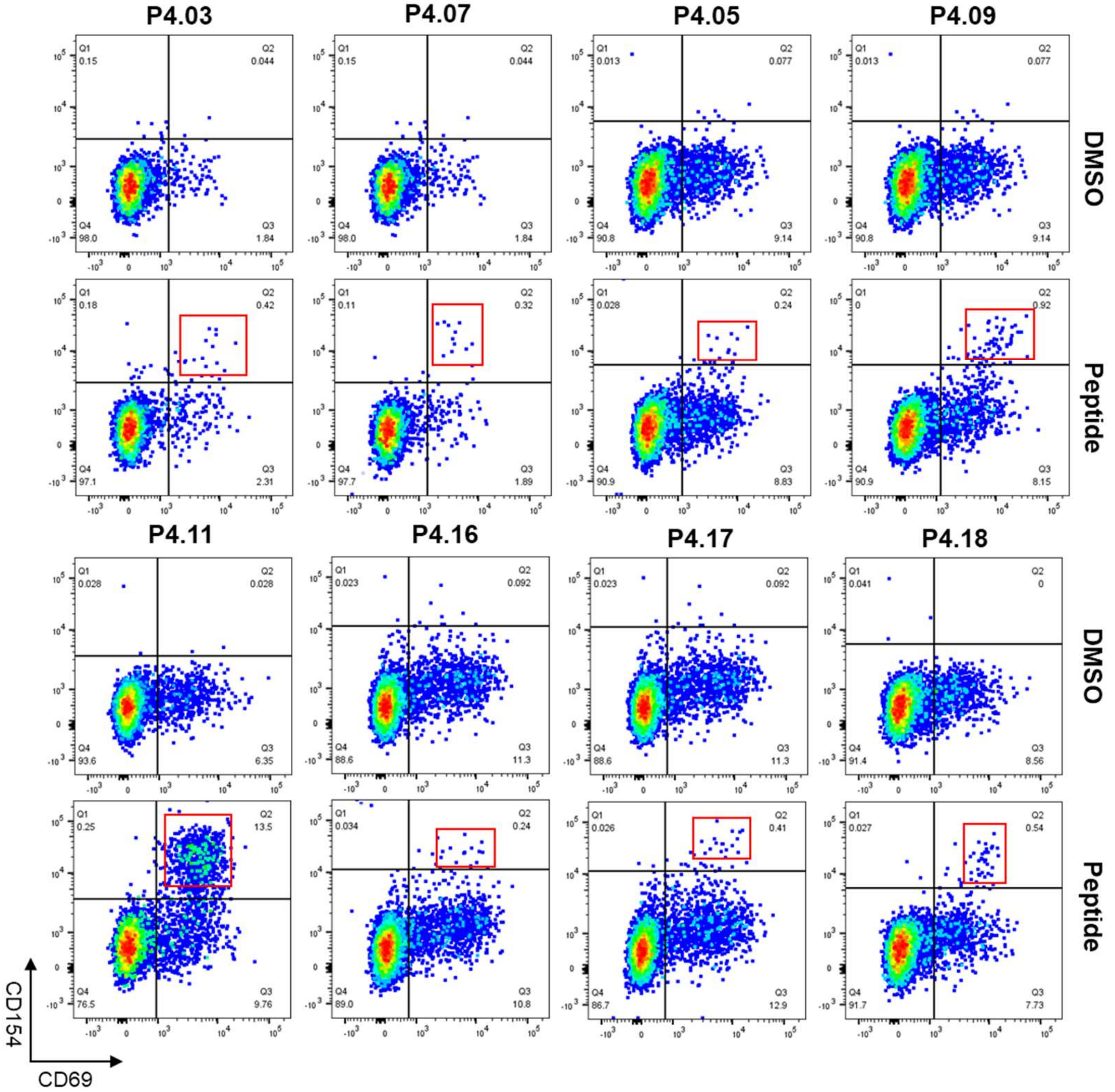


**ESM Fig. 9. Representative FACS plots of positive CD154 AIM responses for each HLA-DRB1*04:01 peptide**. Eight epitopes (P4.03, P4.05, P4.07, P4.09, P4.11, P4.16, P4.17 and P4.18) induced activation of T cells in at least one donor. DMSO was used as negative control.

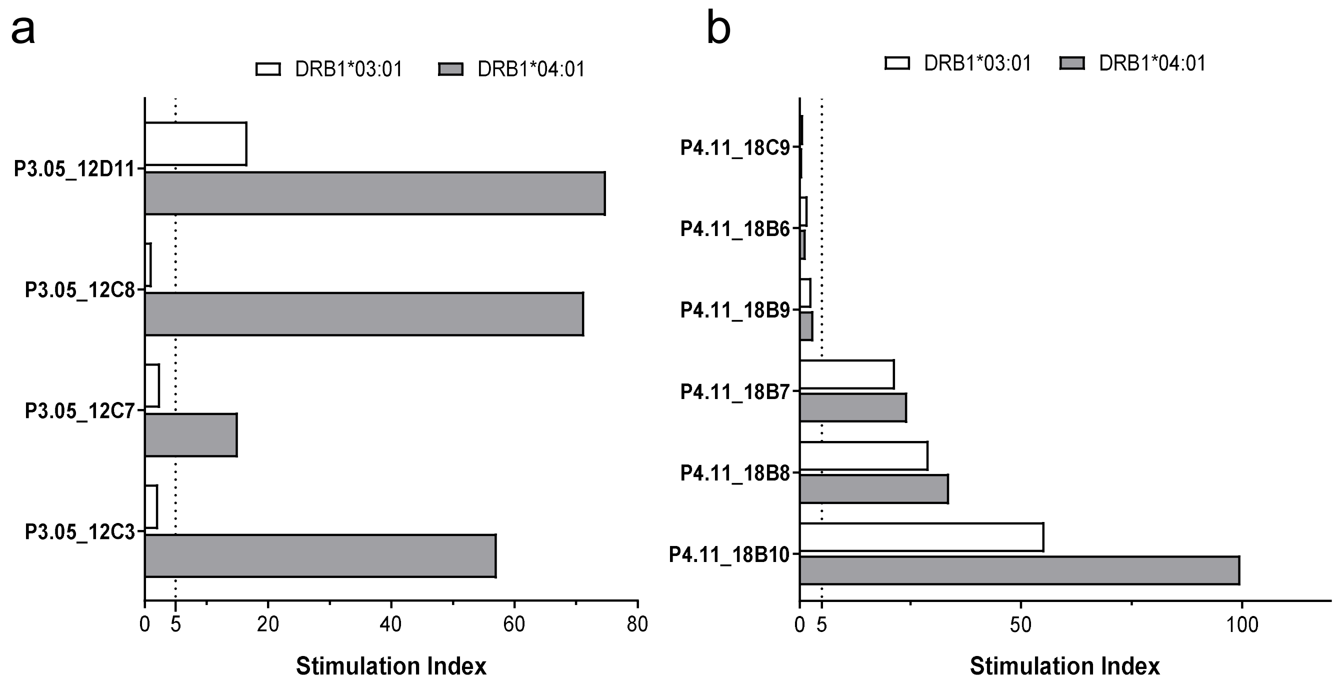


**ESM Fig. 10. Partial overlap between suspected HLA-DR3 and HLA-DR4 peptide presentation.** Stimulation indexes obtained for P3.05 lines expanded from donor 12 (a) and P4.11 lines expanded from donor 18 (b).

**ESM Table 1. List and sequences of primers used for gene expression and m$^6$A analysis by qPCR.**

| Expression analysis | Forward | Reverse |
|---|---|---|
| *UXT-AS1* | TTTGTTTGATGCTCTCTCTCCA | CTCACAGAAGGACAGGGCAT |
| *RAPGEF4-AS1* | TGTTTCAAGGTCACCAAAAGCT | CAGAGGGGAGGGTATTGCTT |
| *ENS00000227066* | CCTACGTTTGGACTGCCCTT | ACATGGCTGTGGTGAATCCA |
| *RPLP0* | CACTGGCAACATTGCGGA | GCAGCATCTACAACCCTGAAG |
| *Lnc13* | AAGGATCATTGCAGGGTCTC | GTGGCCAAAAGAAGTCTGAGTC |

| m$^6$A analysis | Forward | Reverse |
|---|---|---|
| *UXT-AS1* | CTCTCCCTGCTTGAAGTCCC | TCTCGGTCCAGGCATTGTAG |
| *RAPGEF4-AS1* | CAATGGGGCTGAATGATGCA | TCCTGTTTCCTGTCTCCCAC |
| *ENS00000227066_m$^6$A_7* | TTCTGGGCATCAATCACAGC | CTGTCCTCCCTCTGCCTAGA |
| *ENS00000227066_m$^6$A_11-15* | GGAAACTCATCGTGGGACTC | ACACACAAGTCCCTTTTAAACAG |

### ESM Table 2. Sequences of lncRNA candidate ORFs and their potential translations as identified by MS.

| lncRNA gene | Transcript | ORF sequence | ORF translation | Peptide length (aa) |
|---|---|---|---|---|
| *ENSG00000227066* | *ENST00000431027.1* | ATGCATACCAAAGGACATCATTGGCTAACAAATGGTA GACTAACTAAGTACCAAAGCTTGCTCCGTGAAAATCC CTGCATAACCATTGAAGTTTGCAACACCTTGAACTCCA CCACCTTACTCCCAGTGTCAGAGAGCCCAGTCGAACG TAACTATGTAGAGGTGTTGGACACAGTTTATTCTAGCA GGCCTGACCTCCAAGACCATCCTTGGACATCAGTAGA CTGGGAGCTGTACATGGACAGGAGCAGCTTCGTCAAC CCACAAGGAGAGAGGTGTGCAGGATATGCAGTGGTA ACCCTGGATGCTGTCACTGAAGCCAAATCATTGCCCC AGGATACTTCAGCCCAAAAGGTCAAACTCATTGCTTTA ATTTGGGCCTTAGAGCTAAGTGAAGGTAAGACTGTAA ATATTTATACTGACTCTCGGTATGCCTTCTTAACCCTC CAAGTGCATGGGGCATTATATAAAGAAAAAGGCCCAT TGAAATCTGGGGGAAAAGACATAAAGTACCAGCAAGA GATCCTGCAGTTAGTAGAGGCAGTATGGAAGCCCCAA AAGGTGGCAGTCATGCATTGCAGAGGACACCAGCGA GCTTCCACCTCGATTGCCTTGGGGAACTCCCAAGCTG ACTCAGAGGCTTGA | MHTKGHHWLTNGRLTKYQSLLRENPCITI EVCNTLNSTTLLPVSESPVERNYVEVLDT VYSSRPDLQDHPWTSVDWELYMDRSSF VNPQGERCAGYAVVTLDAVTEAKSLPQD TSAQKVKLIALIWALELSEGKTVNIYTDSR YAFLTLQVHGALYKEKGPLKSGGKDIKYQ QEILQLVEAVWKPQKVAVMHCRGHQRAS TSIALGNSQADSEA | 214 |
| *UXT-AS1* | *ENST00000669654.1* | ATGCACCTGCAGGGATCCTCCCTTGTGGCCCTCTCCC TGCTTGAAGTCCCACCCGAAAAGCTTGGGGACATAGC AGCAGCTGCCTCGGTACAGAGGACTAATAAATTTTGTT TGATGCTCTCTCTCCATACTACAATGCCTGGACCGAG ATTCCAGCTTTGGCCATGGGGAGCTCCTTCATGCTGG TTCCTGCATCCTTCAGACATGCCCTGTCCTTCTGTGAG CACTGTCTTACTTTCTGACCCCACAAGATATTTCAGGC TCATCTTGTAG | MHLQGSSLVALSLLEVPPEKLGDIAAAAS VQRTNKFCLMLSLHTTMPGPRFQLWPW GAPSCWFLHPSDMPCPSVSTVLLSDPTR YFRLIL | 90 |
| *RAPGEF4-AS1* | *ENST00000435328.1* | ATGAGAAGAGTTCCCTCTGTCTGCTCTTTCCTCTGCCT GATGACCCAGAGGCTTTGCAGCTGTGATCCTGGCAGC CCCTGGCCACCTCAGGGAGCTGCAGAATTCAAACCCT GGAAATGGAGCTCTGCTTGCAGCTGCACCTGGGGCC TGACCTGCTTAGGACAGGCCTGTCTGGACCTAAGAGA ACCCCACTGCCACCCACTCAGCTACTCCCTAACAGAC TGCAGAGACCAAGTCTTTTTCATTTCTTTATATCCATTG ATTTCTCGAGTTATTGAAACTCAATATTTACTGAGCAC CTACAATGTGCCAGGCACTGTGATAGGCACTGAAGAA AGGGCAAAGGACGAGCAGATAAATTTCCTACTTGCAG GGAACACCGTGGTAGTGAGTAAAATGAGACATTGCAA ATAA | MRRVPSVCSFLCLMTQRLCSCDPGSPW PPQGAAEFKPWKWSSACSCTWGLTCLG QACLDLREPHCHPLSYSLTDCRDQVFFIS LYPLISRVIETQYLLSTYNVPGTVIGTEERA KDEQINFLLAGNTVVVSKMRHCK | 137 |

**ESM Table 3. Summary of in vitro assays and T1D donor characteristics.**

| Subset | Donor | *DRB1* genotype | Sex | Age | T1D duration |
|---|---|---|---|---|---|
| T1D DRB1*03:01 | *1* | *0301/*0401 | male | 38 | 22.1 |
| | *2* | *0301/*0401 | male | 13 | 9.2 |
| | *3* | *0301/*1201 | male | 49 | 47.4 |
| | *4* | *0301/*0404 | male | 21 | 8.7 |
| | *5* | *0301/*1601 | male | 24 | 2.1 |
| | *6* | *0301/*0301 | male | 24 | 11.0 |
| | *7* | *0301/*1302 | female | 19 | 6.4 |
| | *8* | *0301/1310*1302 | female | 16 | 3.7 |
| | *9* | *0301 / *0404 | female | 19 | 2.5 |
| | *10* | *0301/*0901 | male | 14 | 4.9 |
| | *11* | *0301/*1302 | female | 14 | 1.1 |
| | *12* | *0301 / *0401 | female | 51 | 39.3 |
| T1D DRB1*04:01 | *13* | *0401/unknown | female | 56 | 9.6 |
| | *14* | *0401/*0401 | female | 41 | 11.3 |
| | *15* | *0401/*1302 | male | 31 | 6.2 |
| | *16* | *0401/unknown | female | 33 | 7.8 |
| | *17* | *0401/*03 | female | 42 | 11.1 |
| | *18* | *0401/*13 | female | 35 | 8.3 |
| | *19* | *0401/*03 | male | 34 | 7.1 |

**ESM Table 4. Summary of the autoreactive HLA-DRB1*03:01 CD4 T cell lines included in the proliferation assays**. In bold those lines which exceed the SI threshold to be considered as restricted. First two digits of the line label indicate the original donor (eg P3.01 10C9 is line C9 from donor 10 against peptide P3.01).

| Peptide | Line | DRB1*03:01 SI | Peptide | Line | DRB1*03:01 SI |
|---|---|---|---|---|---|
| P3.01 | **10C9** | **54.5** | P3.09 | 10D9 | 1.4 |
| | **10C7** | **11.9** | | 12F11* | 1.2 (3.1) |
| | 10C6 | 2.6 | | 10C5 | 1.2 |
| | 10C8 | 1.3 | | 10C10 | 0.9 |
| P3.02 | **11D10** | **19.6** | P3.11 | **11F9** | **204** |
| | **11D11** | **12.2** | | **11E7** | **77.5** |
| | **11D5** | **7.3** | | **11F4** | **54.2** |
| | 11D3 | 1.3 | | **11E8** | **34.2** |
| | 10B2 | 1.2 | | **12C7** | **18.5** |
| P3.05 | **12D11*** | **16.7 (74.9)** | | **12C8** | **16.8** |
| | **12C7*** | 2.5 (**15.2**) | | **12B8** | **10.4** |
| | **12C3*** | 2.2 (**57.1**) | | **12B4** | **6.7** |
| | 11C3 | 2.1 | | **12C10** | **6.2** |
| | 11C4 | 1.4 | | 10F7 | 3.1 |
| | 11C6 | 1.2 | | 10G9 | 2.7 |
| | **12C8*** | 1.1 (**71.3**) | | 10G8 | 2.2 |
| | 11D6 | 1 | | 11E11 | 2.1 |
| | 11D11 | 0.7 | | 10G5 | 2 |
| P3.09 | **10D4** | **25.9** | | 11F6 | 1.3 |
| | 10C7 | 4.3 | P3.12 | 11B4 | 2.1 |
| | **12E5*** | 1.8 (**11.3**) | | 11C10 | 1.9 |
| | **12F10*** | 1.7 (**22.8**) | | 11B7 | 1.2 |
| | 10D7 | 1.5 | | | |

*HLA-DRB1*04:01-restricted B cell proliferation assays stimulation indexes are shown between parentheses for these donor 12 lines, as the recognized peptide substantially overlaps another DRB1*04:01 peptide

**ESM Table 5. Summary of the autoreactive HLA DRB1*04:01 $CD4^+$ T cell lines included in the proliferation assays**. In bold those lines which exceed the SI threshold to be considered as restricted. First two digits of the line label indicate the original donor (eg P4.05 18B7 is line B7 from donor 18 against peptide P4.05).

| Peptide | Line | DRB1*04:01 SI | Peptide | Line | DRB1*04:01 SI |
|---|---|---|---|---|---|
| P4.05 | **18D2** | **310.6** | P4.16 | 17B3 | 3.4 |
| | **18C9** | **289.7** | | 17B7 | 2.5 |
| | **18C6** | **251.3** | P4.17 | **19B7** | **262.7** |
| | **18C11** | **220.1** | | **17E3** | **197.2** |
| | **18B7** | **213.5** | | **19B10** | **192.8** |
| | **18B2** | **131** | | **17E5** | **132.7** |
| | **14B7** | **79.7** | | **19D2** | **131.6** |
| | **14B10** | **69.3** | | **19B3** | **67.6** |
| | **14B4** | **60.6** | | **17E9** | **32.1** |
| | **14B2** | **51.6** | | **19B9** | **24.1** |
| P4.09 | **14E8** | **51.1** | P4.18 | **15B9** | **315.5** |
| | **14E10** | **45.8** | | **15B6** | **128.4** |
| | **14D4** | **16.2** | | **15D11** | **107.2** |
| | **14D10** | **5.4** | | **15B5** | **54.7** |
| | 18E4 | 2.5 | | **14G11** | **49** |
| P4.11 | **18B10*** | **99.6 (55.4)** | | **14G7** | **18.5** |
| | **18B8*** | **33.7 (29.1)** | | **14G9** | **17.3** |
| | **18B7*** | **24.3 (21.6)** | | **15C11** | **13.9** |
| | 18B9* | 3.1 (2.7) | | **15C10** | **13** |
| | 18B6* | 1.4 (1.8) | | 14E5 | 6.3 |
| | 18C9* | 0.6 (0.7) | | | |

*HLA-DRB1*03:01-restricted B cell proliferation assays stimulation indexes are shown between parentheses for these donor 18 lines, as P4.11 peptide substantially overlaps P3.05.
